# Illuminating the renal response to pH stress with single-nucleus RNA sequencing

**DOI:** 10.64898/2026.01.28.702357

**Authors:** Jianxiang Xue, Krystin Eaton, Omar Rachidi Alaoui, Olga Ponomarova, Kathryn J Brayer, Nathan A. Zaidman

## Abstract

Maintenance of whole-body pH is essential for human health. The kidneys play a crucial role in defending pH homeostasis by excreting excess acid in the urine and returning alkali buffers to the blood. Consequently, renal insufficiency causes serious and harmful effects on pH balance. While a serious and common complication of chronic kidney disease (CKD), pH imbalances themselves appear to be catalysts of kidney injury. Renal adaptations to pH imbalances contribute to compensated acid-base disorders and are vital to correcting whole-body pH. However, overstimulation of these adaptive processes can cause renal inflammation and lead to long-term kidney injury. Surprisingly, the acute and chronic effects of pH challenges on the whole kidney are poorly defined. The upregulation of ammoniagenesis in the proximal tubule due to acidosis, and the coordinated secretion of protons from the collecting ducts is a well-documented phenomenon. However, there is a significant gap in knowledge regarding how the other segments of the nephron respond to acidosis or alkalosis. Therefore, to determine the cell-specific impact of overt metabolic acidosis and alkalosis on the kidney, we performed single-nucleus RNA sequencing on male and female WT mice following 48-hours of acid-base challenge (280mM NH4Cl (acid), 280mM NaHCO3 (alkali), 280mM NaCl (isosmotic control)). The results of our studies reveal the sex-specific single-cell transcriptional response by the kidney to pH imbalances, including a proximal straight tubule cell cluster that arises *de novo* following both acidosis and alkalosis. We label these proximal tubule cells PT S3a and demonstrate that their transcriptional profile is distinct from other injured PT cells that arise from ischemic injury. These studies lay the foundation for future research into the long-term renal adaptations to pH challenges that may lead to renal insufficiency and the development of CKD.

## Introduction

Maintenance of pH homeostasis is essential to human health^1^. Acid-base disorders cause severe morbidities, including bone disorders^2^, skeletal muscle atrophy^3^, electrolyte imbalance^4^, cardiac arrhythmias^5, 6^, failure to thrive in adolescents^7^, and coma. Kidneys are essential regulators of acid-base status^8^. Accordingly, metabolic acidosis is a common complication of chronic kidney disease (CKD)^9–11^. The kidneys maintain pH balance through the reclamation of filtered bicarbonate and the excretion of excess acid or base equivalents in the urine. Most excess acid is carried by phosphate or ammonia buffers, while citrate is the primary base equivalent excreted in the urine. The consumption of a protein-rich western diet increases catabolism of sulfur-containing amino acids (*eg*, methionine, cysteine) leading to net production of 50-70 mEq of H^+^ per day and significantly greater reliance on renal processes to maintain pH homeostasis^12–14^.

In response to metabolic acidosis, the kidneys quickly adapt their metabolism and transport properties to generate new bicarbonate through the catabolism of glutamine^15^. This process, called ammoniagenesis^16^, regenerates bicarbonate ions and partially corrects the acidification of the extracellular fluid by buffering free H^+^. Ammoniagenesis occurs predominantly in the proximal convoluted tubule (PCT) and the adaptive responses in the PCT have been documented in humans^17^ and rodents^15, 18–20^. Beyond the PCT, bicarbonate regeneration requires synchronized transport of NH_4_^+^ throughout the renal tubules^21–24^ and interstitium^25–27^ and ultimately depends on acid-secretion from A-type intercalated cells^28, 29^ in the collecting ducts to finally consolidate bicarbonate. Thus, it is truly the whole kidney that adapts during metabolic acidosis. However, the precise, and cell-specific, transcriptional response of the kidney to an acidosis, or an alkalosis, has not been reported.

Current next-generation transcriptomics techniques offer unique windows of observation into the etiology of pathologies^30–33^, the effects of therapeutics^34, 35^ or physiologically relevant adaptive processes^36^ that induce cell-autonomous responses. The cellular heterogeneity of the kidney makes these techniques especially valuable in understanding renal physiology and pathophysiology^37^. In the present work, we aimed to develop a comprehensive and cell-specific transcriptomic map of the mouse kidney following acute acid-base disturbances. Our goals were three-fold: 1) produce another high-quality single-nucleus transcriptional dataset of the male and female mouse kidney as a public resource, 2) identify transcripts that demonstrate cell-specific sensitivity to pH status, and 3) determine the acute pathological effects of standard rodent acid-base challenges on the kidney. Here, we present a single-nucleus analysis of male and female mouse kidneys following 48-hours of metabolic acidosis or alkalosis. Our results reveal adaptive processes throughout the kidney, confirming decades of research, while adding a trove of new cell-specific data. Furthermore, we report the identification of a subset of proximal straight tubule (PST) cells that arise in both female and male mice during pH imbalances and are transcriptionally distinct from severely injured proximal tubule cells observed in ischemia-reperfusion injury (IRI) models. We propose that tracking this population of PST cells may reveal novel insights into renal repair mechanisms following physiologically relevant challenges.

## Materials and Methods

### Animals, Induction of pH Imbalances, Urine Collection and Blood Analysis

All animal procedures were conducted in accordance with guidelines set forth by the National Institutes of Health *Guide for the Care and Use of Laboratory Animals* and were approved by the University of New Mexico Animal Care and Use Committee. Experiments were performed on 14-week-old male and female C57BL/6J mice (Strain #000664). All mice were acclimated to Techniplast mouse metabolic cages for one week before the start of the experiment. Mice were provided with 0.5% sucrose water *ad libitum* for 48-hour baseline collections. Mice were provided with 280 mM NH_4_Cl, 280 mM NaHCO_3_ or 280 mM NaCl dissolved in 0.5% sucrose water *ad libitum* for 48-hour experimental collections. 24-hour urine samples were collected under mineral oil, centrifuged to remove debris and transferred to a clean Eppendorf tube. Urine pH was measured by a Thermo Scientific Orion Star A214 pH meter with an Orion semi-micro tipped glass pH electrode. Urine electrolytes (Na^+^, K^+^) were measured on a Sherwood Scientific Model 425 Flame Photometer diluted 1:2000 in 100ppm Li^+^ standard and operated with non-odorized propane. Urine creatinine was measured with the Creatinine Reagent Set (Horiba Instruments, C7539-625) diluted 1:25 in accordance with manufacturer’s instructions. Urine NH_3_ was measured with the fluorescent Ammonia Assay Kit (Abcam, ab272538) diluted 1:750. Creatinine and ammonia measurements were performed on a BioTek Synergy HT microplate reader. At the conclusion of the 48-hour experiment, retro-orbital blood was collected by micro-hematocrit capillary tubes directly into BD Microtrainer Lithium Heparin blood collection tubes and immediately analyzed on an Abbot i-Stat1 handheld blood analyzer using CG8^+^ cartridges.

### Nuclei Isolation and snRNA-seq

Mice were euthanized by cervical dislocation in accordance with IACUC protocols. Kidneys were quickly excised, decapsulated, cut in half and immediately frozen in liquid nitrogen. Kidneys were stored at −80 C for three weeks. Nuclei were isolated from a single half-kidney in 1 mL Lysis Buffer containing (mM in DEPC treated water): 250 Sucrose, 50 Citric Acid. DEPC (diethyl pyrocarbonate) was added to Lysis Buffer at 1:600 dilution immediately before use. Tissue was lysed on ice in a Dounce homogenizer, then serial filtered through a 70 µm and 35 µm nylon mesh screen to remove debris. Nuclei were pelleted then washed in 1mL Wash Buffer containing (mM in DEPC treated water): 250 Sucrose, 50 Citric Acid, 20 DTT, 1% BSA. 10000 U Ambion RNase Inhibitor (Invitrogen, 40 U/µL) was added to 50 mL Wash Buffer immediately before use. Nuclei were filtered through a 20 µm nylon mesh screen and fixed for 24-hours in Fixation Buffer (10x Genomics) and then quenched with Quenching Buffer (10x Genomics). cDNA library construction was performed as recommended by 10X Genomics and sequenced on a Singular Genomics G4. Sample data was demultiplexed and fastq files were generated onboard the G4. Fastq files were aligned and genes/cells were counted using CellRanger (v7.1, 10X Genomics)^38^ against the mouse reference genome (mm10) provided with CellRanger. Seurat (v5.0.1), an R package (v4.3.2), was used for downstream analysis^39, 40^. Cells (nuclei) expressing fewer than 250 genes, or more than 19000 genes, and with mitochondrial gene expression content greater than 2% were removed. Genes expressed in fewer than 10 cells were also removed. Data were transformed using SCTransform as implemented in Seurat^41^ with mitochondrial genes and the difference between G2M and S cell cycle phase scores regressed out. Data from all samples were integrated using Seurat’s standard integration workflow. Principle component (PC) analysis and an elbow plot were used to visualize the variance and select PCs for unsupervised clustering. Clusters were determined using the FindNeighbors and FindClusters function with default parameters and a sufficiently high-resolution parameter to capture biological variability. Doublet cells were removed manually based on markers, and any cell expressing multiple cell-type markers was removed. Differential gene expression was determined using the ‘wilcox’ method as implemented in Seurat.

### Pseudo-Time Analysis, AddModuleScore, and Pathway Analysis

Pseudotime analysis was performed with Monocle3^42^ (v1.3.7) as implemented in Seurat. Briefly, subsets of all PT cells, or treatment specific cells, were imported (as.cell_data_set), preprocessed and clustered. The Seurat cell embeddings were transferred to the Monocle object to maintain UMAP consistency. After clustering, learn_graph (use_partition = FALSE) was used to calculate trajectories and order_cells to calculate the psuedotime, with the root manually set to “PT S1”. Each cell in the proximal tubule clusters was scored for expression levels of genes in each gene module (Supplemental Tables) using AddModuleScore in Seurat. Resulting UMAPs used min and max cutoffs of 0 and 1.5, respectively. Pathway analysis of differentially expressed genes were tested for pathway enrichment using clusterProfiler^43^ (v4.14.6) against the Molecular Signatures Database MSigDB^44–47^ (msigdb, v1.14.0).

### Metabolite extraction, derivatization, and GC-MS analysis

Frozen kidney tissues (20–70 mg of tissue per sample) were homogenized in 1 ml of 80% methanol using acid washed glass beads. FastPrep-24 bead beater was run for 1 min at 6 m/s, then samples were cooled on ice for 1 min. Homogenization-cooling cycle was repeated for a total of five times. Next, samples we incubated on a nutating mixer for 15 min at room temperature. Extracts were centrifuged at 13000 rpm for 10 minutes to clear debris. 200 µl of extract supernatant were transferred into glass inserts and dried in a SpeedVac concentrator. Dried extracts were derivatized in a two-step process. First, samples were incubated with 20 µL of 20 mg/mL methoxyamine hydrochloride in anhydrous pyridine at 37°C for 60 minutes. Then, 50 µL of *N*-methyl-*N*-(trimethylsilyl)trifluoroacetamide (MSTFA) was added, and samples were incubated at 37°C for 3 hours. After an additional 5 hours of room temperature incubation, samples were analyzed using Agilent 8890 gas chromatograph coupled to a 5977C mass selective detector as previously described^48^. One microliter of derivatized sample was injected in splitless mode onto a HP-5ms capillary column (30 m × 0.25 mm, 0.25 μm film thickness) with helium as the carrier gas at a constant flow of 1.0 mL/min. The GC oven temperature was programmed as follows: initial temperature 80°C (held for 2 minutes), ramped at 5°C/min to 280°C, and held for 1 minute. The injector temperature was 230°C. Mass spectrometry was conducted in electron impact (EI) ionization mode at 70 eV with a source temperature of 230°C and quadrupole temperature of 150°C. The MS was operated in scan mode (m/z 30–500), with a solvent delay of 5 minutes. Metabolite identification and relative quantification were performed using Agilent MassHunter software by matching retention times and fragmentation patterns to both in-house and NIST libraries. Peak areas were normalized to total ion count.

### Statistics and Data Availability

Comparisons between treatment groups were restricted to each sex. All metabolic parameters were subject to two-way ANOVA followed by Tukey (Fig. 1, Sup. Fig. 1) or Holm-Sidak (Sup. Figs. 2, 4, 5, 7) multiple comparison tests. Statistical analysis of snRNAseq data was performed Seurat.

**Figure 1:**
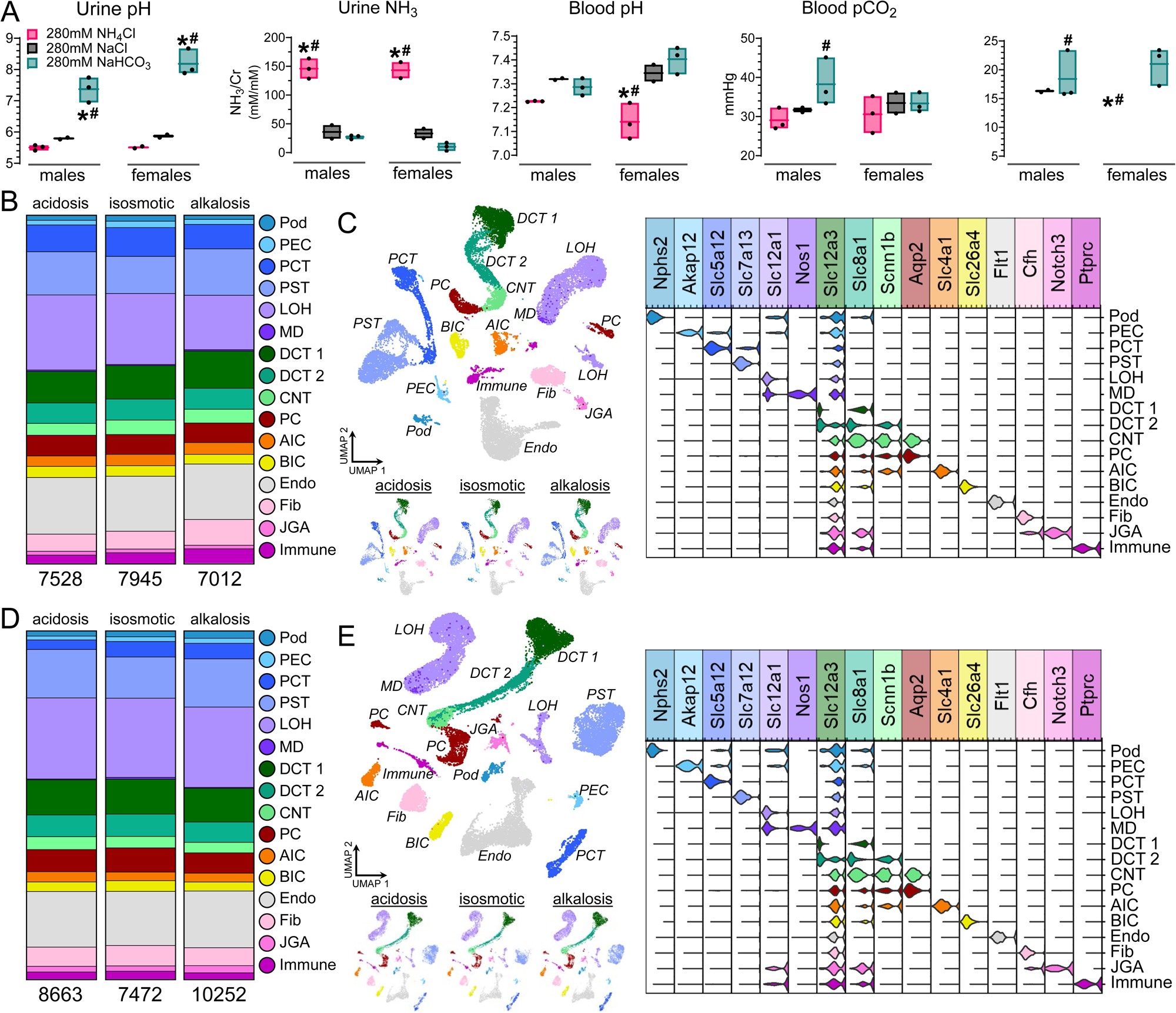
Single-nucleus RNA sequencing following induction of metabolic acidosis and alkalosis identified 16 cell-clusters. A) Measurement of urinary pH and ammonia, and blood pH, pCO2 and bicarbonate, demonstrate induction of metabolic acidosis and alkalosis by addition of solute to the drinking water. B) Distribution of cell-types identified by snRNAseq in male mice. C) UMAP and violin-plot of all male cells combined identified 16 distinct cell-clusters. Inset shows UMAPs by condition. D) Distribution of cell-types identified by snRNAseq in female mice. C) UMAP and violin-plot of all female cells combined identified 16 distinct cell-clusters. Inset shows UMAPs by condition. * padj<0.05 test vs control, # padj<0.05 acidosis vs alkalosis.

## Results

### Induction of metabolic acidosis or alkalosis

To determine the transcriptional effects of acute acid-base disturbance on the kidney, we induced a metabolic acidosis or alkalosis in 14-week-old male and female C57Bl/6 mice for 48 hours by addition of 280mM NH_4_Cl or NaHCO_3_ to the drinking water. To control for greater salt consumption, isosmotic control groups were given 280mM NaCl in the drinking water for 48 hours. Urine pH was significantly higher in male and female NaHCO_3_ treated animals compared to isosmotic controls (Figure 1A; male: 7.37±0.4 vs. 5.79±0.04 p_adj_=0.0004, female: 8.18±0.4 vs. 5.87±0.06 p_adj_<0.0001). Though urine pH was not significantly different in male or female NH_4_Cl treated animals compared to controls (Figure 1A; male: 5.50±0.09 vs. 5.79±0.04 p_adj_=0.51, female: 5.51±0.04 vs. 5.87±0.06 p_adj_=0.43), urinary ammonia was significantly increased (Figure 1A; male: 145.93±17.4 vs. 35.99±16.5 p_adj_<0.0001; female: 143.31±19.7 vs. 33.32±11.8 p_adj_<0.0001). Male NH_4_Cl treated mice demonstrated trends towards lower blood pH and HCO_3_^-^ compared to isosmotic controls, while female NH_4_Cl treated mice had significantly reduced blood pH and HCO_3_^-^ compared to controls. 24-hour water intake, urine flow and changes in body mass are presented in Supplemental Figure 1. Additionally, urinary K^+^ was significantly elevated in female NH_4_Cl treated mice compared to baseline (Supplemental Figure 1; 520.3±52.9 mEq/L vs. 217.9±140.2 mEq/L, p_adj_=0.0037) and trended higher in males (317.3±31.9 mEq/L vs. 217.3±51.2 mEq/L, p_adj_=0.096). Urinary Na^+^ was significantly increased from baseline in both male and female NaCl and NaHCO_3_ treated animals (Supplemental Figure 1). Altogether, these results demonstrate that acute acid-base imbalances were induced in male and female mice.

### Identification of cell-type specific effects to acute acid-base disturbance

After 48-hour inductions, mice were euthanized, and kidneys were harvested for snRNA-seq. Following quality-control measures to remove low-quality nuclei or doublet GEMs, a total of 7528 nuclei from male NH_4_Cl treated mice, 7945 nuclei from male isosmotic control mice, and 7012 nuclei from male NaHCO_3_ treated mice were analyzed (Figure 1B). Similarly, a total of 8663 nuclei from female NH_4_Cl treated mice, 7472 nuclei from female isosmotic control mice, and 10252 nuclei from female NaHCO_3_ treated mice were analyzed (Figure 1D). Unsupervised clustering identified 16 distinct cell-clusters in both male (Figure 1C) and female kidneys (Figure 1E), including podocytes (Pod), parietal epithelial cells (PEC) and fibroblasts (Fib).

To increase the cellular resolution of the transcriptional effects of acid-base disturbance, we further refined the nuclei into 27 and 29 clusters for males (Figure 2A) and females (Figure 2G), respectively. The proximal convoluted tubule (PCT, Figure 1) was subdivided into three distinct clusters: PT S1 labeled by high Slc5a12 and Slc5a2 (male) or Akr1c21 (female) expression, PT S2 labeled by high Slc13a3 and low Slc5a12 expression, and PT S1-2 labeled by high Slc5a12 and Slc13a3 expression. The proximal straight tubule (PST, Figure 1), labeled by Slc7a13 or Slc7a12 for males and females, respectively, was also subdivided into two distinct clusters called PT S3a and PT S3b. In male mice, a single cluster marked by Bst1, and having detectable Slc14a2 expression, was identified that corresponded with the thin descending and ascending limbs of the loop of Henle (DTL-ATL). In contrast, three Bst1^+^ DTL-ATL clusters were identified in female mice: DTL-ATL 1 labeled by high Bst1 and low Slc4a11^49^ expression, DTL-ATL 2 labeled by high Slc4a11 expression, and DTL-ATL 3 labeled by high Slc14a2 expression. The thick ascending limb of the loop of Henle was subdivided into three clusters for both male and female mice: mTAL (medullary) labeled by high Clcnka expression in the males and high Nccrp1 expression in the females, cTAL 1 (cortical) labeled by high Slc12a1 and low Car15 expression in the males and high Slc12a1 and low Casr expression in the females, and cTAL 2 labeled by high Car15 and low Clcnka expression in the males and high Casr and low Nccrp1 expression in the females. A cluster of nuclei with high Nos1 expression corresponding to macula densa cells (MD) was also identified in male and female mice.

**Figure 2:**
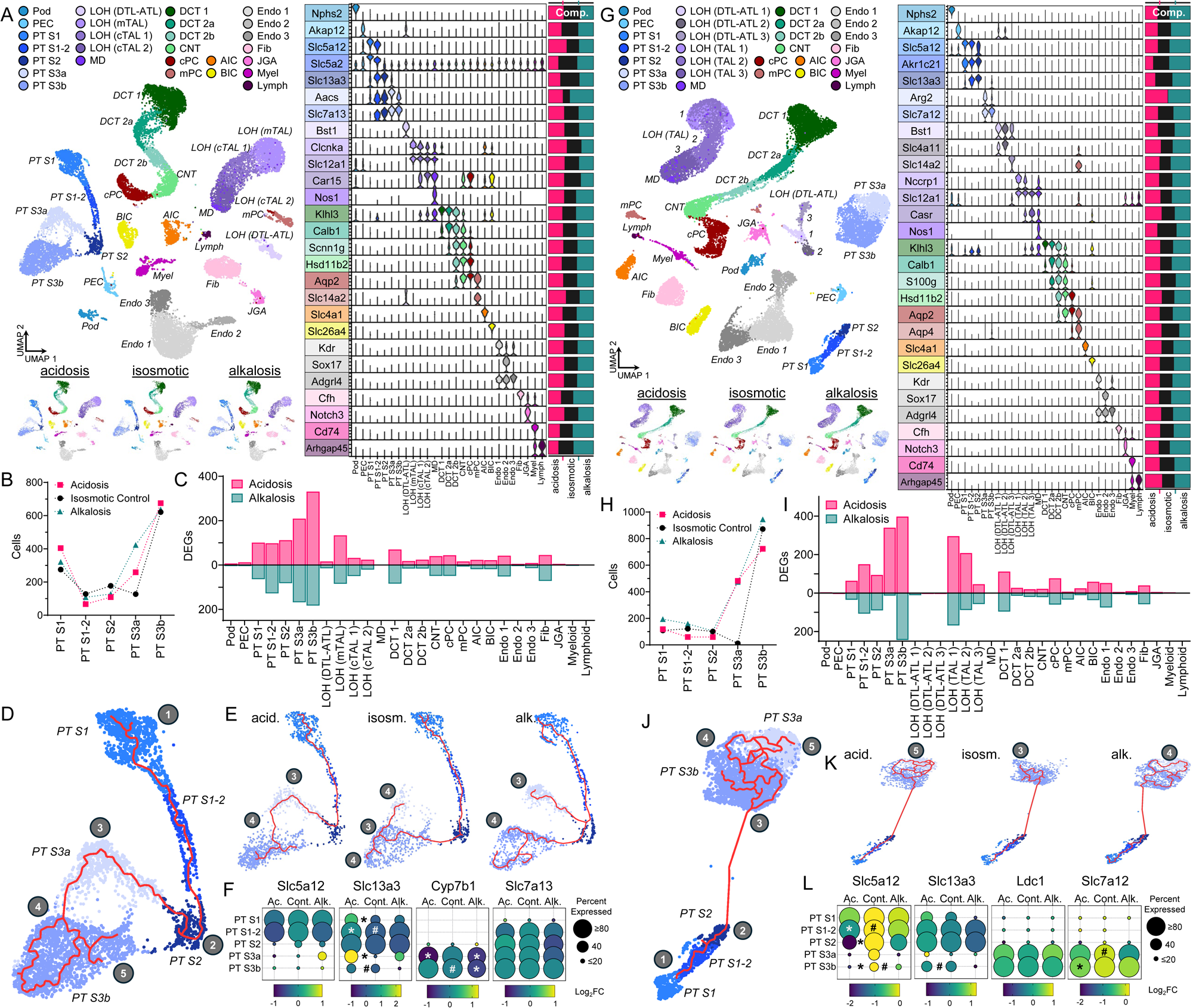
Refinement of clusters reveals a subset of proximal tubule cells that are susceptible to acid/base disturbance. A) UMAP and violin-plot of refined male cells identifying 27 transcriptionally distinct clusters. Inset shows UMAPs by condition. Percent composition (treatment) of each cluster presented to the right of violin-plot. B) Total number of cells by treatment in each of the five proximal tubule clusters. C) Total number of differentially expressed genes compared to isosmotic control in each of the 27 male clusters. D,E) Pseudo-time analysis of all cells combined (D) and subdivided by treatment (E) for the five proximal tubule clusters. Earliest time = 1, latest time = 5. F) Dot plots of proximal tubule segment genetic markers presented by treatment. G) UMAP and violin-plot of refined female cells identifying 29 transcriptionally distinct clusters. H) Total number of cells by treatment in each of the five proximal tubule clusters. I) Total number of differentially expressed genes compared to isosmotic control in each of the 29 female clusters. J,K) Pseudo-time analysis of all cells combined (J) and subdivided by treatment (K) for the five proximal tubule clusters. Earliest time = 1, latest time = 5. L) Dot plots of proximal tubule segment genetic markers presented by treatment. * p_adj_<0.05 test vs control, # p_adj_<0.05 acidosis vs alkalosis.

Cells belonging to three distinct distal convoluted tubule phenotypes (DCT 1, 2a, 2b) were identified as previously reported in male and female kidneys^50^. Additionally, collecting duct principal cells were subdivided into cortical and medullary (Slc14a2^+^) populations in both sexes. Endothelial cells were subdivided into three Adgrl4^+^ clusters in both male and female mice: Endo 1 labeled by high Kdr (VEGF receptor 2) expression, Endo 2 labeled by high Sox17 and low Kdr expression, and Endo 3 labeled by low Kdr, low Sox17 and high Adgrl4 expression. The immune cell cluster was subdivided into nuclei belonging to either Cd74^+^ myeloid derived cells or Arhgap45^+^ lymphoid derived cells.

The proportion of cells originating from acidosis, isosmotic control, or alkalosis treatment groups was determined for each cluster (Figures 1A, G). Generally, each of the three treatment groups accounted for roughly 1/3 of the nuclei in each cluster. However, male and female PT S3a clusters were notably populated by cells belonging to either NH_4_Cl or NaHCO_3_ treated mice (Figures 1B, H). Strikingly, only 12 female nuclei from the isosmotic control group sorted into PT S3a, strongly suggesting that PT S3a cells are a *de novo* population of PST cells arising from pH imbalance.

To determine the cell-type specific effects of metabolic acidosis or alkalosis, we performed differential gene expression (DEG) analysis where we compared all treatment groups to each other (for each sex independently). Figures 1C and 1I show the total number of significant DEGs in each cluster for NH_4_Cl or NaHCO_3_ treated mice compared to isosmotic controls. A majority of DEGs were detected in the proximal tubules in both sexes, with acidosis PT S3a and S3b having the greatest number of DEGs.

To further understand the relationship of PT S3a to the other proximal tubule cell clusters, we performed pseudo-time analysis to determine where PT S3a cells reside along the trajectory of proximal tubule cell differentiation (Figures 1D, J). Early pseudo-time, corresponding to the initiation of the trajectory, corresponds to PT S1 and then progresses through the “later” PCT clusters. Analysis on all male nuclei revealed PT S3a’s trajectory to reside between the PCT and PT S3b. Analysis by treatment confirmed this result in males (Figure 1E), suggesting that PT S3a are dedifferentiated PT S3 cells. Pseudo-time analysis on female proximal tubule clusters offered a different result than the males. Analysis on all cells combined, as well as by treatment, demonstrated that PT S3a resides “later” along the trajectory than PT S3b in females. Gene expression analysis of proximal tubule markers confirmed that PT S3a nuclei possess an expression profile that most closely aligns with PT S3 in both male and female mice. These results confirm that PT S3a is a *de novo* cluster of cells that arise in the distal proximal tubule following acute acid-base disturbance.

### PT S3a is a subset of injured cells in the PST

To better understand the origin and function of the PT S3a cluster, we tested the hypothesis that these cells were recruited for bicarbonate reclamation, though it would be unclear why NaHCO_3_ treated mice would respond in that manner. Nonetheless, we performed a scoring experiment using the “AddModuleScore” function in Seurat, which amalgamates the expression of the 14 genes from the KEGG Pathway: *Proximal Tubule Bicarbonate Reclamation* (ko04964; Figures 3A, 4A). Quantification of the score shows significantly greater values in all PT clusters during metabolic acidosis compared to control but not metabolic alkalosis (Supplemental Figures 2, 5). PT S3a was significantly different from control and alkalosis in the acidosis male mice. Female acidosis PT S3a had a significantly different score only compared to alkalosis. The ammoniagenesis scores were significantly greater in all acidosis treated PCT clusters compared to controls and alkalosis, as well as greater than acidosis PST clusters, confirming the primary localization of ammoniagenesis to the PCT. Moreover, significantly greater scores in the acidosis PT S3b clusters reveals that all proximal tubule segments are recruited, to some extent, to participate in bicarbonate reclamation during acute metabolic acidosis.

**Figure 3:**
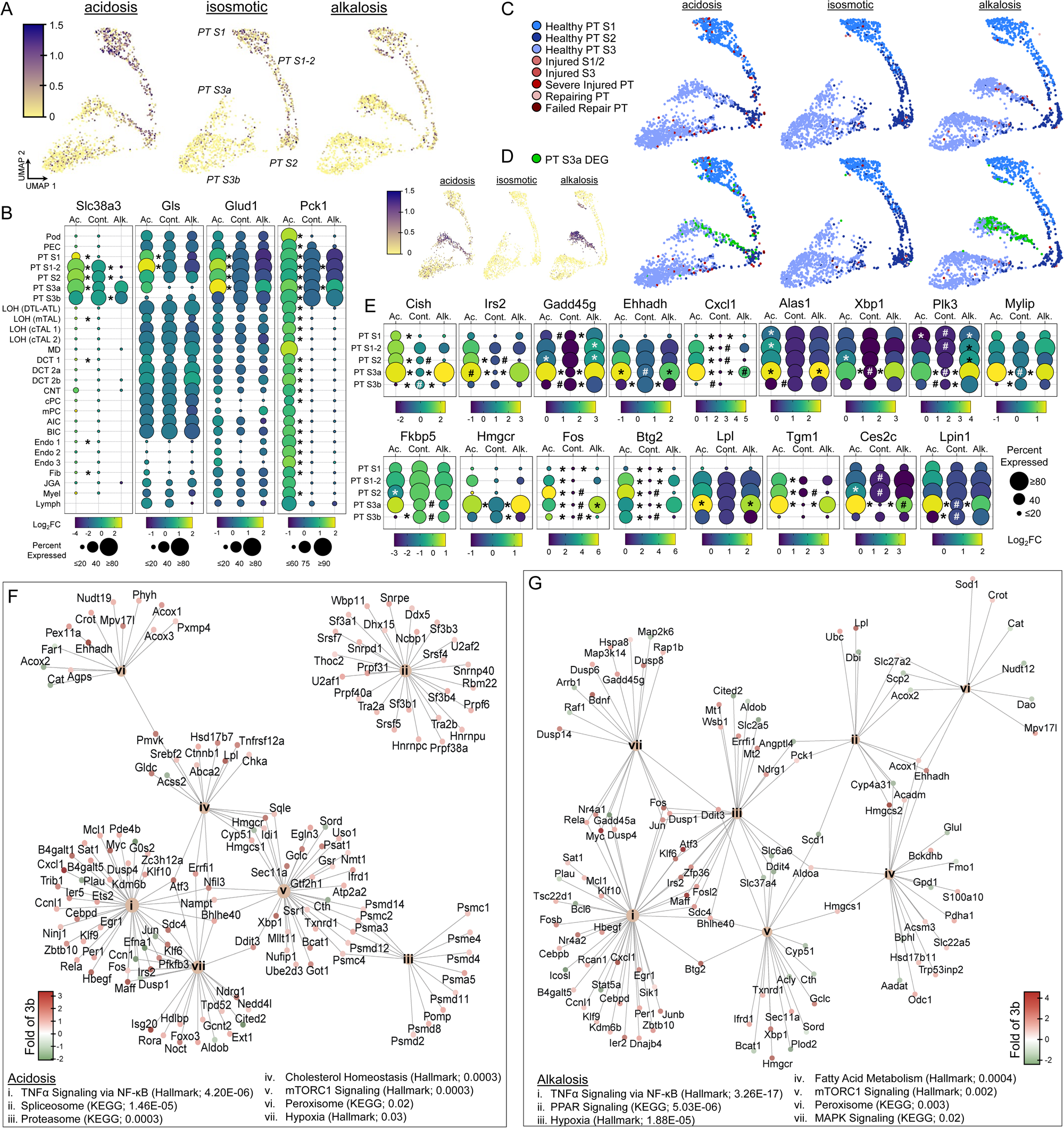
Male PTS3a is a subset of PST cells that experience a mild injury during acute acid/base disturbance. A) Ammoniagenesis score expression by treatment for the five proximal tubule clusters. Darker color indicates higher score. B) Dot plots of transcripts critical to ammoniagenesis for all 27 male cell clusters presented by treatment. C) Winning score expression by treatment for the five male proximal tubule clusters. Scores were compiled for eight proximal tubule profiles. The profile with the highest score was declared the “winner”. D) Winning score expression following the introduction of PTS3a DEG profile. Inset shows PTS3a score expression by treatment for the proximal tubule clusters (purple-yellow). E) Dot plots for the most enriched genes in PT S3a compared to S3b. F,G) Pathway analysis for PTS3a DEGs compared to PTS3b during acidosis (F) and alkalosis (G) showing the top seven enriched pathways from KEGG and Hallmark datasets. Data is presented as a gene-concept network (cnetplot). Gene color (red-green) indicates PT S3a fold change compared to PT S3b. p_adj_ included in parentheses. * p_adj_<0.05 test vs control, # p_adj_<0.05 acidosis vs alkalosis.

To further validate our results, we analyzed four key transcripts critical to proximal tubule ammoniagenesis/bicarbonate reclamation to assess the effects of acute metabolic acidosis or alkalosis on their expression (Figures 3B, 4B). Slc38a3, the transcript for the basolateral glutamine transporter SNAT3, is significantly upregulated in the proximal tubule during metabolic acidosis and significantly downregulated during alkalosis. Slc38a3 is minimally expressed in PT S1 in isosmotic controls and is undetected in PT S1 in NaHCO_3_ treated mice. In contrast, a subset of PT S1 cells upregulate Slc38a3 during metabolic acidosis, demonstrating the nephron’s capacity to recruit additional cells to participate in bicarbonate reclamation. Gls, the transcript for kidney-type glutaminase, which catalyzes the hydrolysis of Glutamine to Glutamate + NH_4_^+^, is upregulated in the PCT clusters (males: PT S1, S1-2; females: PT S1-2, S2) and has notably low expression in S3 clusters in the male isosmotic controls. Though transcription is not significantly upregulated, 41% of PT S3a cells express Gls in the male acidosis group (17% isosmotic control, 18% alkalosis), supporting the possibility that these cells were recruited to participate in ammoniagenesis. Glud1, the transcript for glutamate dehydrogenase, which catalyzes the oxidative deamination of Glutamate to 2-oxoglutarte + NH_4_^+^, and Pck1, the transcript for phosphoenolpyruvate carboxykinase (PEPCK), which converts oxaloacetic acid into phosphoenolphyruvate that either supplies the TCA cycle with pyruvate (via pyruvate kinase) or is consumed for gluconeogenesis, are both upregulated in the proximal tubule during metabolic acidosis. Pck1 is upregulated throughout the nephron in the male acidosis group, whereas Pck1 is downregulated along the nephron in the female alkalosis group owing to the notably high percent of all cells expressing Pck1 in the isosmotic controls. These results demonstrate that the proximal tubule generally upregulates these transcripts during an acidosis, while an alkalosis causes either downregulation (Slc38a3, Pck1) or no change (Gls, Glud1). Furthermore, the transcriptional data partially supports the hypothesis that in the male acidosis mice PT S3a are PST cells that are recruited to participate in ammoniagenesis by the induction of Gls.

While our analysis of ammoniagenesis revealed some clues about the function of PT S3a cells, the scoring data from the males (Figure 3A) strongly suggested that PT S3a cells were not simply newly recruited ammoniagenic tubule cells. To assess whether PT S3a were injured cells, we similarly employed the “AddModuleScore” function in Seurat using transcriptional profiles of healthy and ischemia-reperfusion injury (IRI) proximal tubule cells^30^ (Figures 3C, 4C). In this analysis the gene list with the highest amalgamated score was labeled the winner for each nuclei, allowing for unbiased identification of “healthy” or “injured” cells. Analysis of the results showed that nearly all nuclei were most associated with transcriptional profiles aligning with “healthy” PT segments (Supplemental Figures 3, 4). The most severely injured cells were dispersed in the PT clusters, but not PT S3a. The average AddModule Score for all injury gene lists were very low. The results of this analysis demonstrate that if PT S3a cells were experiencing an injury it was transcriptionally distinct from an IRI.

Next, we compiled a list of the top 15 DEGs from PT S3a and then performed another “AddModuleScore” analysis to determine if the PT S3a genotype could be discerned from the other PT transcriptional profiles (Figures 3D, 4D). Predictably, our curated gene list identified many of the PT S3a nuclei in the challenged male and female mice, demonstrating that PT S3a cells have a transcriptional profile that is distinct from healthy PT S3 cells as well as severely injured, repairing or failed repairing PT cells (Supplemental Figures 3, 4).

To further identify the transcriptional profile of PT S3a, we performed pairwise differential gene expression analysis between PT S3a and PT S3b from male and female treatment groups. Although PT S3b had many DEGs based on our analysis in Figures 2C and 2I (by cluster, to controls), the cluster was present in all three male and female treatment groups and was estimated to be composed of “healthy” PT S3 cells (Figures 3D, 4D). Thus, our analysis asked the question “what are the distinguishing transcriptional differences between PT S3a and PT S3b?” The most enriched genes based on log_2_ fold-change are presented in Figures 3E and 4E. Notably, most of the enriched transcripts are present in PT S3a independent of the variety of acid-base challenges, which aligns with the finding that the PT S3a cluster develops in both the acidosis and alkalosis UMAPs. There were also transcriptional differences between male and female PT S3a cells. For example, Cxcl1 was enriched in male PT S3a but not female, while Arg2 was enriched in female PT S3a but not male (Supplemental Figure 8).

**Figure 4:**
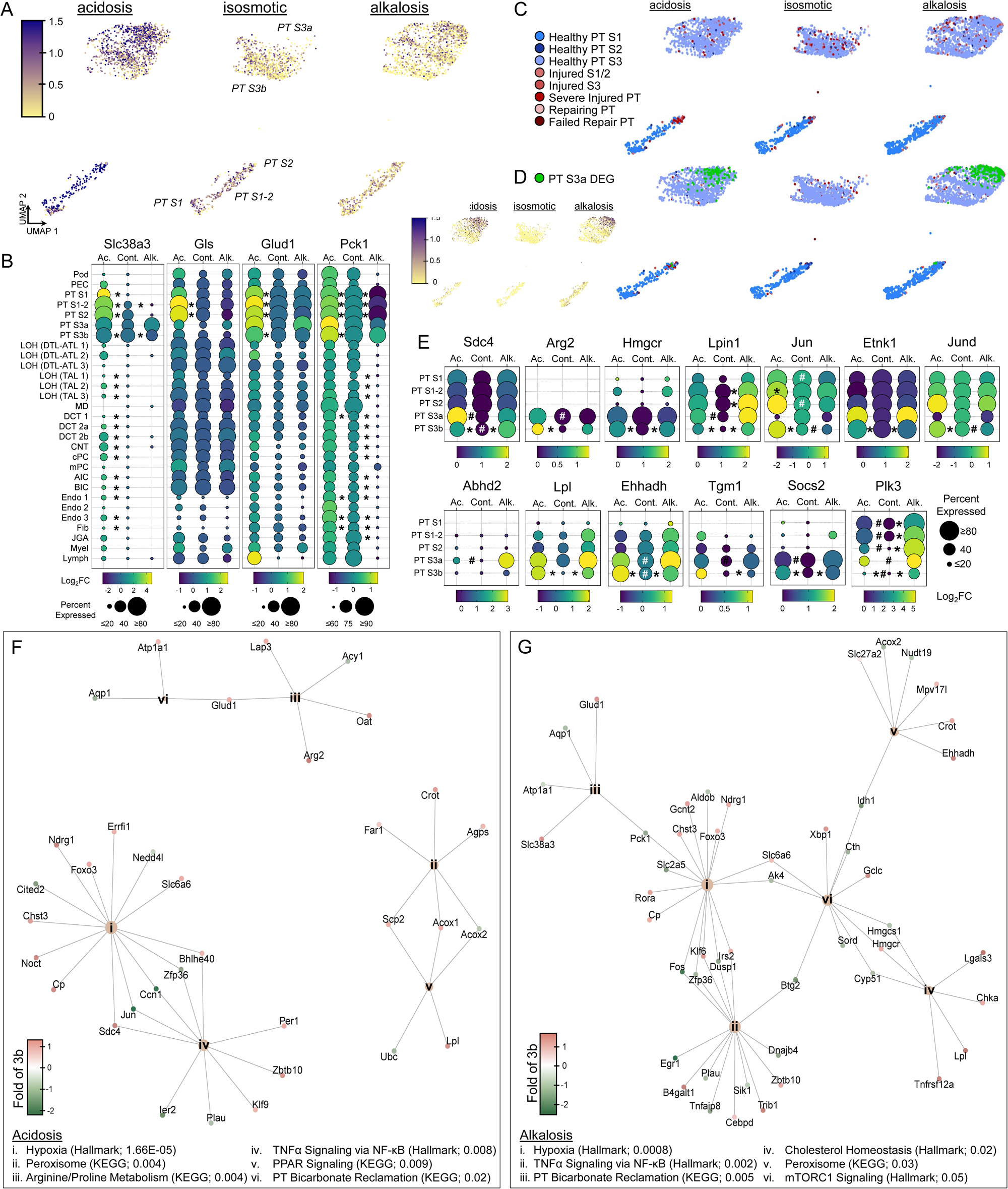
Female PTS3a are a subset of PST cells that experience an injury during acute acid/base disturbance and are not present at baseline. A) Ammoniagenesis score expression by treatment for the five proximal tubule clusters. Darker color indicates higher score. B) Dot plots of transcripts critical to ammoniagenesis for all 29 female cell clusters presented by treatment. C) Winning score expression by treatment for the five female proximal tubule clusters. Scores were compiled for eight proximal tubule profiles. The profile with the highest score was declared the “winner”. D) Winning score expression following the introduction of PTS3a DEG profile. Inset shows PTS3a score expression by treatment for the proximal tubule clusters (purple-yellow). E) Dot plots for the most enriched genes in PT S3a compared to S3b. F,G) Pathway analysis for PTS3a DEGs compared to PTS3b during acidosis (F) and alkalosis (G) showing the top seven enriched pathways from KEGG and Hallmark datasets. Data is presented as a gene-concept network (cnetplot). Gene color (red-green) indicates PT S3a fold change compared to PT S3b. * p_adj_<0.05 test vs control, # p_adj_<0.05 acidosis vs alkalosis.

Using the pairwise DEG analysis between PT S3a and S3b, we performed pathway analysis using GSEA datasets. The most enriched orthology-mapped Hallmark Pathways and KEGG gene sets are presented as gene-concept networks (CNET plot) in Figures 3F/G and 4F/G. Complete pairwise pathway analysis is included in the Supplemental Tables. The most enriched pathway for both male challenge groups was *TNF*α *Signaling via NF-*κ*B*, suggesting that PT S3a cells had experienced an inflammatory response that triggered an NF-κB-mediated transcriptional response. Key genes in this pathway include chemokine Cxcl1 and c-Myc. Other enriched pathways in the male mice include *mTORC1 Signaling*, *Spliceosome*, and *Peroxisome*, suggesting that the male PT S3a cells are injured cells mounting recoveries through increased transcription and translation. Notably, gene signatures associated with autophagy or apoptosis were not enriched. In the female mice, the most enriched pathway for both challenge groups were *Hypoxia* from the curated Hallmark gene sets. *TNF*α *Signaling via NF-*κ*B* was also significantly enriched in the female PT S3a clusters, though the total number of genes associated with the pathway were fewer than the males. This trend was apparent throughout the gene set enrichment analysis for the female PT S3a clusters. The results of our analysis suggest that cells clustering in PT S3a are mildly injured PST cells that arise *de novo* after acute acid-base disturbances.

### Effects of acid-base disturbance on the whole kidney

Next, we performed pathway analysis on all DEGs between each treatment group and isosmotic controls to better understand the effects of acid-base disturbances on the entire kidney. Enriched Hallmark pathways are presented as circos plots in Figures 5A and 6A. Several enriched pathways were unique to specific clusters as well as type of acid-base disturbance. The pathways with the greatest representation among all clusters were Hypoxia (males: 23 acidosis/16 alkalosis, Figure 5B; females: 8 acidosis/13 alkalosis, Figure 6B) and TNFα Signaling via NFκB (males: 23 acidosis/23 alkalosis, Figure 5C; females 18 acidosis/20 alkalosis, Figure 6C), suggesting that metabolic stress and mild inflammatory responses were pervasive throughout the kidney. Once again, several genes associated with each pathway were only differentially expressed in one treatment group or the other. Five transcripts were differentially expressed in many clusters irrespective of type of acid-base disturbance in both male and female mice, suggesting a role in renal stress response (Figures 5D, 6D). Dusp1 and Zfp36 have been implicated in DKD ferroptosis, while Egr1, c-Fos and Nr4a1 encode transcription factors that are critical to cell proliferation and differentiation.

**Figure 5:**
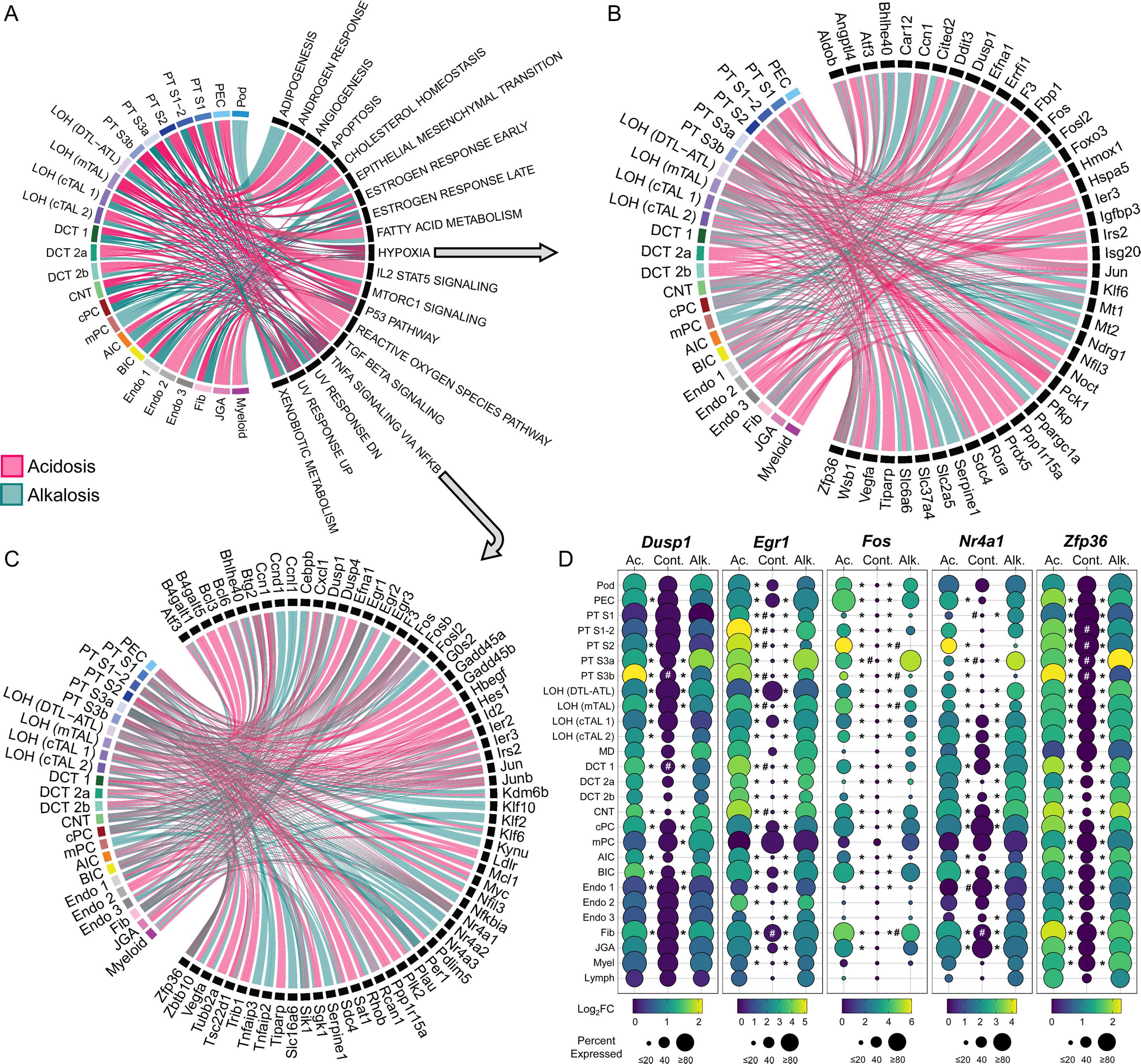
Male gene set enrichment analysis reveals an inflammatory response to acute acid/base disturbance throughout the kidney. A) Circos plot of Hallmark gene set enrichment by treatment and cell cluster. B,C) Circos plot of Hallmark Hypoxia (B) and TNFα-signaling via NFκB (C) pathway showing genes that are differentially expressed by treatment and cell cluster. D) Dot plots of selected genes from (B) and (C) highlighting the differential expression by treatment and throughout the kidney. * p_adj_<0.05 test vs control, # p_adj_<0.05 acidosis vs alkalosis.

**Figure 6:**
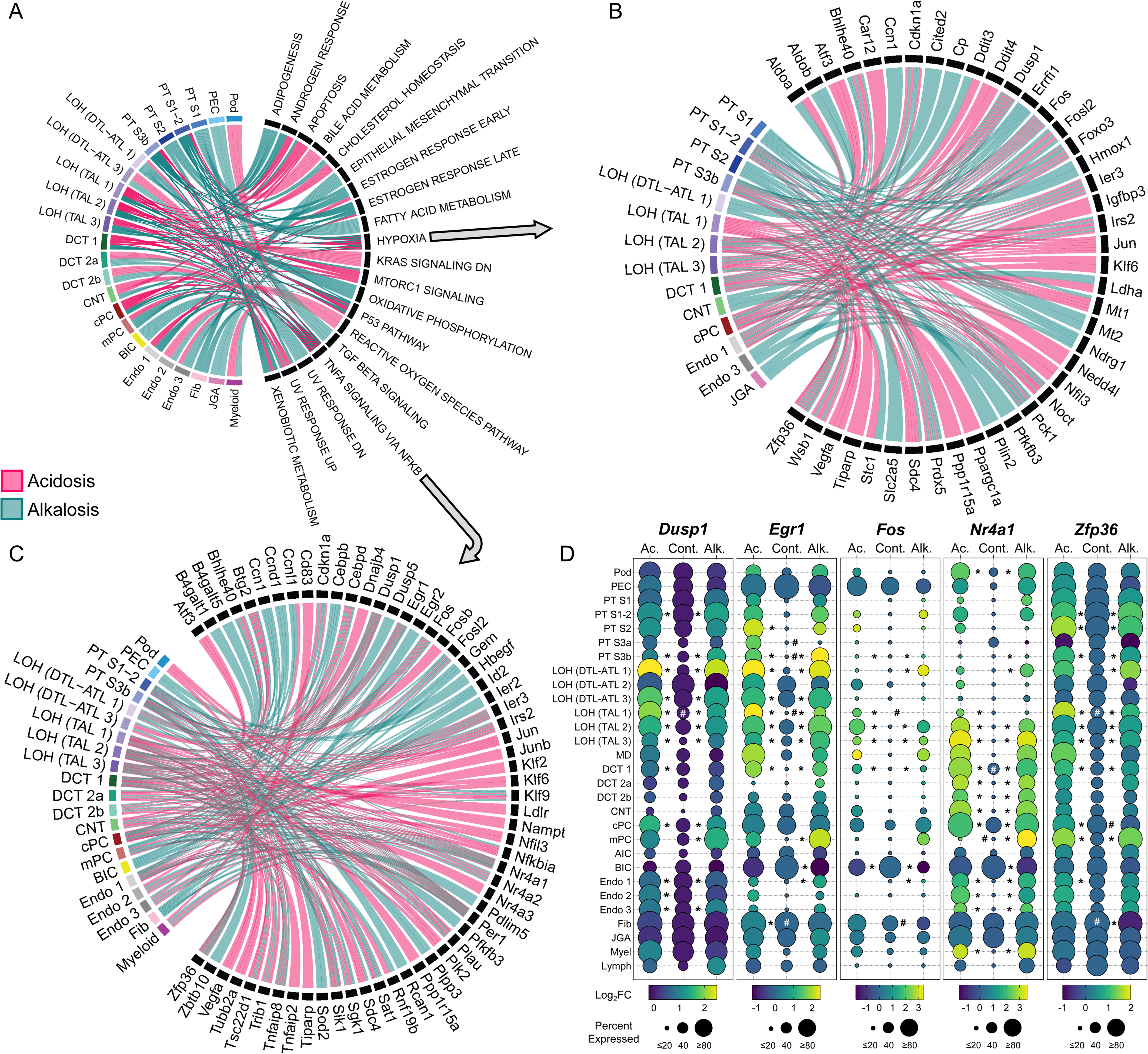
Female gene set enrichment analysis reveals an inflammatory response to acute acid/base disturbance throughout the kidney. A) Circos plot of Hallmark gene set enrichment by treatment and cell cluster. B,C) Circos plot of Hallmark Hypoxia (B) and TNFα-signaling via NFκB (C) pathway showing genes that are differentially expressed by treatment and cell cluster. D) Dot plots of selected genes from (B) and (C) highlighting the differential expression by treatment and throughout the kidney. * p_adj_<0.05 test vs control, # p_adj_<0.05 acidosis vs alkalosis.

Another transcript that was represented in the TNFα Signaling via NFκB circos plot in both sexes was Per1, a circadian rhythm gene, suggesting that circadian control was potentially impacted by the acute acid-base disturbances. Per1 was differentially expressed throughout the kidney in male and female mice (Figure 7A/D), but surprisingly, was more sensitive to NaHCO_3_ treatment in the males. This trend in the male mice was consistent for other period family genes (Per2, Per3) as well as Dbp, a transcription factor under strict circadian control, but was not apparent in female mice. All male mice were euthanized between 0900 and 1100 MDT, and all female mice were euthanized between 1000 and 1130 MDT. Male and female alkalosis mice were the last group of animals euthanized and therefore it is possible that the transcriptional results are impacted by experimental timing (isosmotic controls < acidosis < alkalosis). However, the response in the NaHCO_3_ treated male mice is not observed in the female NaHCO_3_ treated mice, which are paired groups temporally. These results suggest that acute metabolic alkalosis, but not metabolic acidosis, significantly alters circadian rhythms in male kidneys.

**Figure 7:**
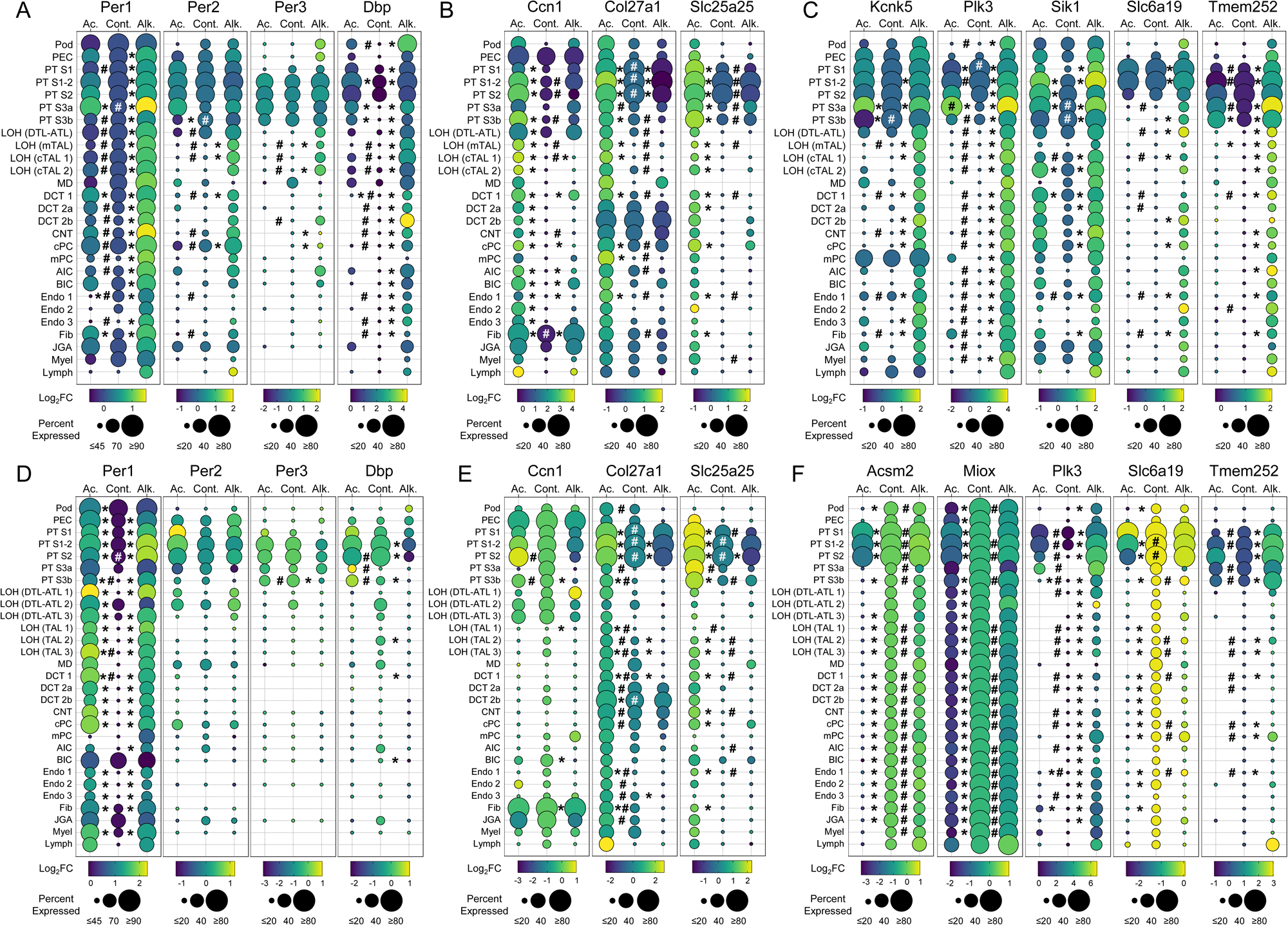
Differential regulation of genes during metabolic acidosis or alkalosis. A,D) Male (A) and female (D) dot plots for select circadian clock genes. B,E) Male (B) and female (E) dot plots for genes differentially regulated during metabolic acidosis. C,F) Male (C) and female (F) dot plots for genes differentially regulated during metabolic alkalosis. * p_adj_<0.05 test vs control, # p_adj_<0.05 acidosis vs alkalosis.

Next, we searched for other transcripts that were predominately affected by either NH_4_Cl or NaHCO_3_ treatment. In the male mice, three transcripts were identified that showed preferential differential expression in the acidosis group: Ccn1, Col27a1, and Slc25a25 (Figure 7B). Both Ccn1 and Col27a1 encode proteins associated with the extracellular matrix, while Slc25a25 encodes a Ca^2+^-sensitive mitochondrial ATP-Mg/P_i_ exchanger that may protect cells from oxidative stress. In the female mice, both Col27a1 and Slc25a25 were preferentially upregulated throughout the kidney after NH_4_Cl treatment, like the males (Figure 7E). We also identified five transcripts that demonstrated preferential upregulation in the NaHCO_3_ treated male mice (Figure 7C). Kcnk5 transcribes the TWIK-related acid-sensitive (TASK-2) two-pore K^+^ channel; Plk3 encodes a kinase that is upregulated during hypoxia- and ischemia-reperfusion AKI models and involved in oxidative stress-induced DNA damage. Sik1 transcribes salt inducible kinase 1 which plays a protective role in the AA-induced AKI-CKD mouse model, Slc6a19 encodes the Na^+^-dependent neutral amino acid transporter B(0)AT1, and Tmem252 encodes a transmembrane protein with no known physiological function. In the female mice, Plk3 followed a similar upregulation during alkalosis as the males, but the trend was much less pronounced for Slc6a19 and Tmem252 (Figure 7F). Two transcripts in female mice were preferentially downregulated in the NH_4_Cl treated mice: Acsm2 and Miox (Figure 7F). Acsm2 transcribes a kidney-specific acyl-CoA synthetase that is downregulated by renal injury and has previously been reported to be restricted to the PT. Miox transcribes *myo*-inositol oxygenase, which supplies precursors that are utilized in the pentose phosphate pathway (generation of NADPH). Miox has also been implicated as a potential biomarker of AKI. A similar trend was observed for Miox in male mice, though Miox was preferentially upregulated in alkalosis compared to controls and acidosis, not simply downregulated in acidosis as observed in the females. Acsm2 transcription had no substantial trend in the male mice, though was robustly detected in all clusters, like the females.

### Transcripts involved with acid-base homeostasis are relatively stable during acute pH imbalance

One of the major objectives of our experiment was to understand how transporters essential to renal acid-base regulation are affected by acute acidosis or alkalosis. Dot plots for selected transcripts are presented in Figure 8 (males 8A, females 8B). In male NH_4_Cl treated mice Car2 was significantly upregulated in AICs, while Slc4a9 (AE4) and Slc26a4 (Pendrin) were significantly downregulated in AICs and BICs, respectively, compared to isosmotic controls. The transcript for the electrogenic sodium bicarbonate cotransporter 1 (Slc4a4) was significantly downregulated in PT S1-2 and S2 in male NH_4_Cl treated mice, as well as PT S1 in male NaHCO_3_ treated animals. Slc12a1, the transcript for the sodium-potassium-chloride cotransporter NKCC2, was significantly downregulated in the LOH (mTAL) cluster in male acidosis mice compared to isosmotic controls and alkalosis. Rhbg transcript was significantly upregulated in the CNT, AIC and BIC clusters in NH_4_Cl treated female mice compared to NaHCO_3_ treated mice. Slc26a4 was significantly upregulated in BICs in the female alkalosis group compared to isosmotic controls and acidosis. Slc12a1 transcript was significantly upregulated in all three female TAL clusters in the alkalosis group compared to the acidosis group. These results largely confirm previous studies on the effects of acidosis or alkalosis on transcripts involved with renal pH homeostasis.

**Figure 8:**
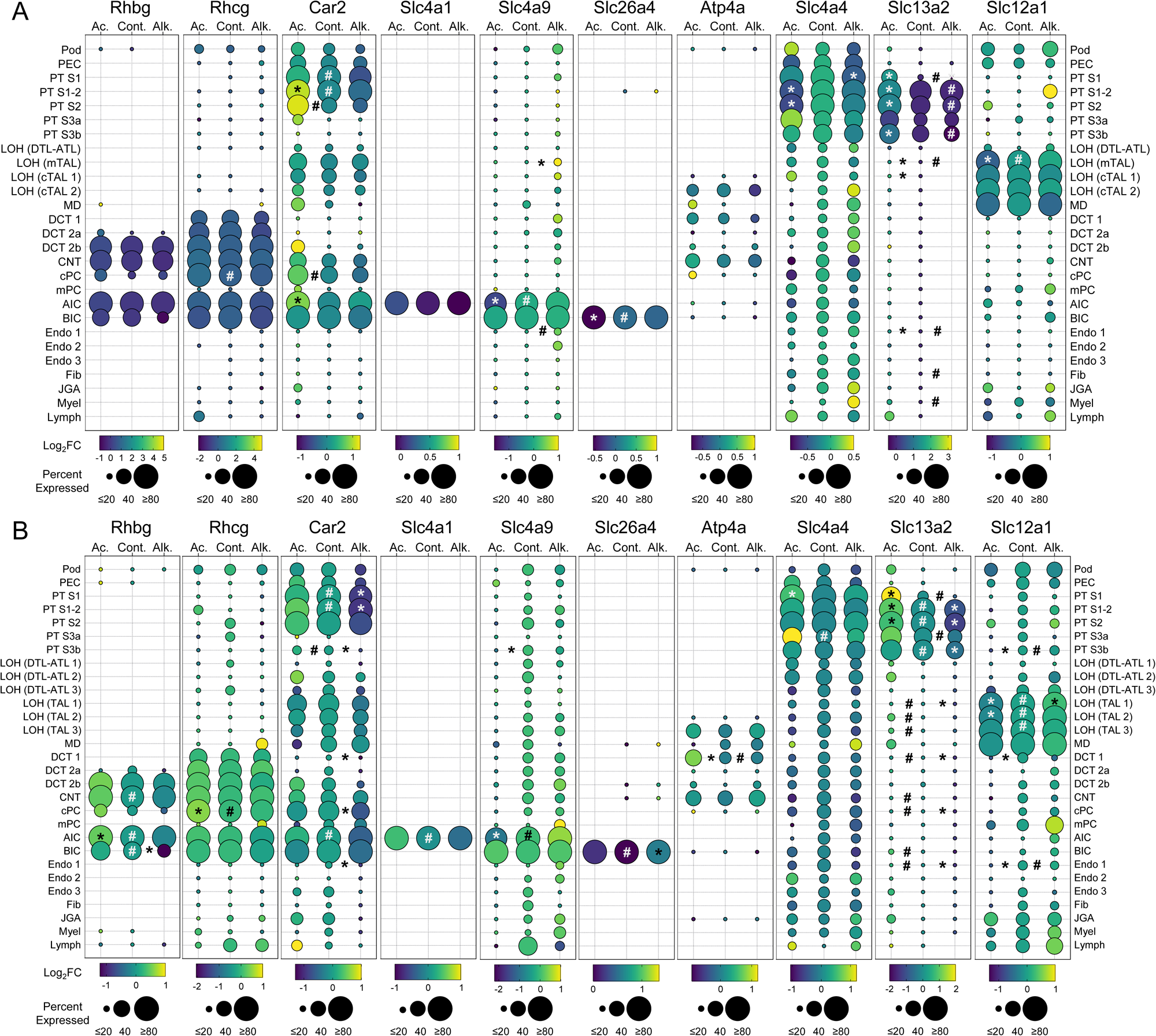
Acute acid/base disturbance has minimal effects on select transcripts involved in pH homeostasis. A,B) Male (A) and female (B) dot plots for genes involved in acid-base homeostasis showing differential expression by treatment. * p_adj_<0.05 test vs control, # p_adj_<0.05 acidosis vs alkalosis.

### Acidosis stimulates the polyamine synthesis pathway in male mice

We noted that Azin1, Odc1, and Sat1 were all significantly upregulated in male PT S3a cells from NH_4_Cl treated mice (Figure 9A). These three transcripts encode critical components of the polyamine synthesis pathway. Odc1 encodes the first and rate-limiting enzyme in polyamine synthesis and has previously been shown to play an essential role in DKD hyperfiltration. While Azin1, Odc1, and Sat1 were differentially expressed in PT S3a in male acidosis mice, there was no effect by either treatment in the female mice (Figure 9B). To determine if polyamines were increased in male kidneys during metabolic acidosis, we performed gas chromatography-mass spectrometry (GC-MS) on matched kidney samples from the snRNAseq experiment. Putrescine, the product of ODC, and spermidine were significantly more abundant in male NH_4_Cl treated mice compared to isosmotic controls as well as NaHCO_3_ treated mice. Spermidine was also increased in male NaHCO_3_ treated mice compared to controls. There were no significant changes in polyamine abundance in the female mice. These results demonstrate that acute metabolic acidosis stimulates renal production of polyamines in a sex-dependent manner.

**Figure 9:**
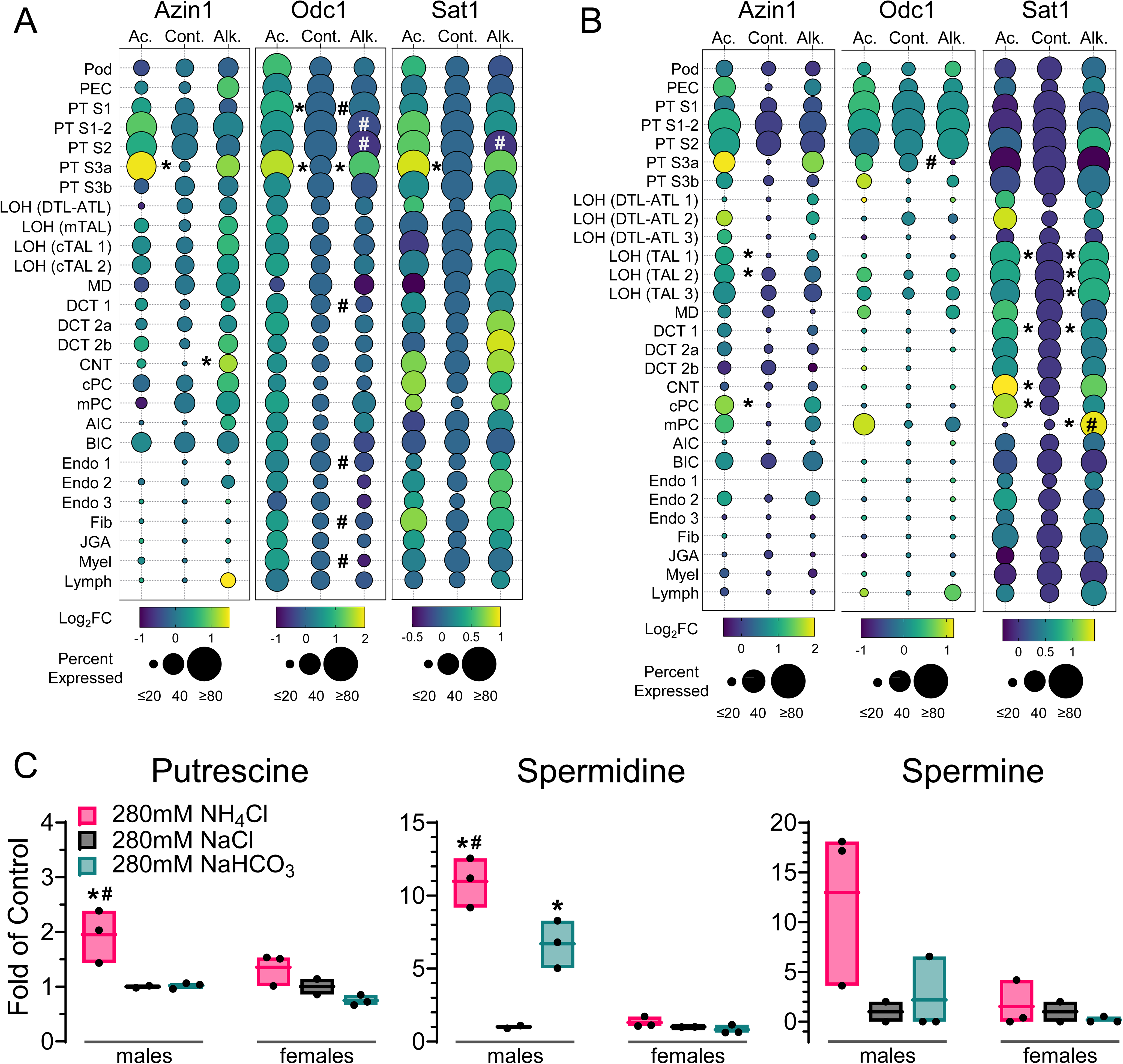
Acute acidosis stimulates putrescine and spermidine synthesis in males but not females. A,B) Dot plots showing male (A) and female (B) transcripts for enzymes involved in polyamine synthesis. C) GC-MS detection of polyamines in snRNAseq-matched whole kidneys. Data is normalized peak area presented as fold of control (280mM NaCl). * p_adj_<0.05 test vs control, # p_adj_<0.05 acidosis vs alkalosis.

## Discussion

The kidneys serve a critical role in maintaining whole body pH homeostasis. In response to an acidosis, the kidneys adapt to increase bicarbonate regeneration and excrete excess acid in the urine. While in response to an alkalosis, the kidney adapts to increase H^+^ retention and excrete excess base units in the urine. Thus, the kidney significantly contributes to compensated acid-base disorders. Though these gross physiological responses are medical canon, the complete cell-specific transcriptional responses to acute pH imbalances in the kidney had not been described. Here, we employed snRNA-seq to illuminate the effects of acute metabolic acidosis or alkalosis on the kidney with single-cell resolution. Our results reveal the renal transcriptional response to acute pH imbalances and highlight the cell-specific adaptability, and susceptibility, of the kidney to this variety of challenges.

Several previous studies assessed transcriptional and translational changes in the mammalian kidney following an acidosis. The most comprehensive of these was conducted by Nowik and colleagues in 2007^18^ and utilized RNA microarrays to determine gene expression regulation in the murine kidney after 48-hours and 7-days 280mM NH_4_Cl treatment. Our work is largely supportive of their findings, including proximal tubule segment-specific expression of ammoniagenesis transcripts such as Slc38a3 and Pck1, significant upregulation of the mitochondrial phosphate carrier Slc25a25 during acidosis, and induction of both adaptive (ammoniagenesis, ATP synthesis) and repair (cell proliferation, differentiation) gene networks. Our study adds significant new information to this seminal study, including analysis of both female and male mice, isosmotic control animals, and the inclusion of NaHCO_3_^-^ challenged mice. Moreover, the resolution of our snRNAseq data reveals the cell-specific expression of transcripts along the proximal tubules and throughout the whole kidney that had previously been limited to micro dissected proximal tubules or bulk kidney analysis. Nonetheless, the results of the Nowik paper withstand the test of time.

Several maladaptive processes have been reported to cause acidosis induced renal injury. One hypothesis argues that increased single-nephron ammoniagenesis in the injured kidney causes interstitial accumulation of ammonia, which leads to tubulointerstitial fibrosis and GFR decline through complement activation^51, 52^. Another hypothesis suggests that acid stress chronically stimulates hormones, such as Angiotensin II and Endothelin-1, that activate renal acid excretion but also advance renal inflammation and fibrosis^53–55^. More recently, Bugarski and colleagues revealed a cell-autonomous mechanism where the proximal tubule accumulates lipids during acute metabolic acidosis. In this model, the stimulation of ammoniagenesis reduces the availability of NAD^+^ for beta oxidation, leading to the formation of large multilamellar bodies and significantly reduced solute uptake. Supplementing NAD or NAM prevented the accumulation of lipids in the PT and restored reabsorption, suggesting that chronic stimulation of ammoniagenesis is incompatible with other PT functions. Furthermore, the accumulation of lipids was determined to preferentially affect PT S2. Our data demonstrates that the late PCT segments (PT S1-2, S2) bear the burden of glutamine uptake from the blood (Slc38a3/SNAT3) and are therefore quite exposed to the high energetic demands of ammoniagenesis. Whether PT S3a represents cells that are also experiencing this type of injury remains to be determined. These cells, we hypothesize, are localized beyond the cortico-medullary boundary and may be very susceptible to changes in energetic demand due to the reduced availability of O_2_ in the renal medulla and their reliance on glycolytic energy production. Thus, PT S3a cells may represent another cellular injury caused by competing energetically demanding functions. Future experiments that determine if NAD or NAM supplementation protects the kidney from pH challenges would be appropriate. Notably, the appearance of PT S3a cells in both the acidosis and alkalosis groups suggests their injury is independent of the direction of pH imbalance.

Another significant finding of our experiments was the stimulation of an inflammatory response throughout the kidney during both acidosis and alkalosis. The upregulation of immediate early genes (IEGs) including Dusp1, Egr1, Fos, Nr4a1 and Zfp36 throughout the kidney demonstrates the responsiveness of the whole kidney to acid-base challenges. Egr1 expression increased the severity of acute kidney injury in both IRI and nephrotoxic mouse models^56^ and was recently linked to FGF2-stimulated fibrosis^57^. Additionally, Dusp1 preserves mitochondrial integrity after AKI^58^ and, together with Zfp36, was implicated as a ferroptosis-related diagnostic marker of DKD^59^. These studies collectively reveal the significance of IEG signaling in renal injury and demonstrate the severity of acute acidosis or alkalosis on the kidney. While the long-term impact of a single pH challenge was not explored in this study, the stimulation of IEGs suggests that there may be lasting effects of acidosis/alkalosis that could contribute to the onset of renal insufficiency.

Metabolic acidosis is a common feature of CKD, and cohort studies have demonstrated an association between metabolic acidosis and CKD progression^60, 61^ and death^62, 63^. In fact, the Chronic Renal Insufficiency Cohort (CRIC) Study reported that the risk of developing ESKD among CKD G2-4 patients was 3% lower per 1.0mEq/L increase in serum bicarbonate^64^. Furthermore, recent studies have demonstrated that subclinical acidosis is a significant risk factor for CKD progression. A retrospective study of three CKD patient cohorts (RENVAS^65^, PUMA^66^, NNRD^67^) reported subclinical acidosis (serum bicarbonate > 22mEq/L) in 67% of all patients using a novel urine acid-base score (urinary pH + [NH_4_^+^])^68^. The study found that H^+^ retention was associated with a larger decrease in GFR after 18 months and significantly higher risk of CKD progression over 6 years. Thus, developing a greater understanding of how acid- or alkali-stress impacts the long-term health of the kidneys may reveal novel therapeutic interventions, or dietary guidelines, that could stem the expanding worldwide CKD epidemic^69, 70^.

While the present study reveals a trove of new information about how the kidney responds transcriptionally to pH challenges, there are many questions that persist. For example, we exclusively studied the effects of acute strong metabolic acidosis or alkalosis. The threshold for a renal transcriptional response, therefore, remains to be understood. Another relevant question is how the kidney transcriptionally responds to chronic low-grade pH challenges, such as subclinical acidosis or a high protein diet. Incorporation of epigenetic analysis and longitudinal studies would reveal how the kidney adapts to pH challenges and expose maladaptive pathways that could be targeted before renal function declines. Another question that arises from our studies is the long-term fate of PT S3a cells in the kidney. Lineage tracing of cycling cells (Ki67^+^) has previously been employed to track proliferating cells after IRI^36^. However, this technique relies on cells entering mitosis, and it is unclear if the pH challenges employed in the present work stimulate cell division. We are currently investigating other transgenic mouse models to trace and isolate PT S3a cells in the kidney after return to pH homeostasis that does not depend on cell division. Importantly, this approach would allow us to assess epigenetic, transcriptional and metabolic changes in these and other cells following a variety of challenges.

In conclusion, we have provided a comprehensive single-cell map of the murine kidney following acute acidosis and alkalosis. We show segment-specific upregulation of ammoniagenesis enzymes as well as the *de novo* formation of PT S3a cells, which appear to be mildly injured cells in the proximal straight tubule. In addition, we highlight sex-differences in the kidney and how the male kidney upregulates synthesis of spermidine in response to pH imbalances. Investigators that are interested in how a specific cell-type responds to pH challenges are invited to query the provided data or to use the cluster specifications to apply to their own single-cell investigations.

## Acknowledgements

This research was partially supported by UNM Comprehensive Cancer Center Suppor Grant NCI P30CA118100 and made use of the Analytical and Translational Genomics shared resources.

## Author Contributions

NAZ conceptualized the study; JX, KE, ORA, and NAZ perfromed the experiments; JX, KE, and NAZ analyzed *in vivo* data; KJB and NAZ analyzed transcriptomic data; OP analyzed GC-MS; NAZ and KJB made figures and drafted the manuscript; all authors approved the final version.

---

**Supplemental Figure 1:**
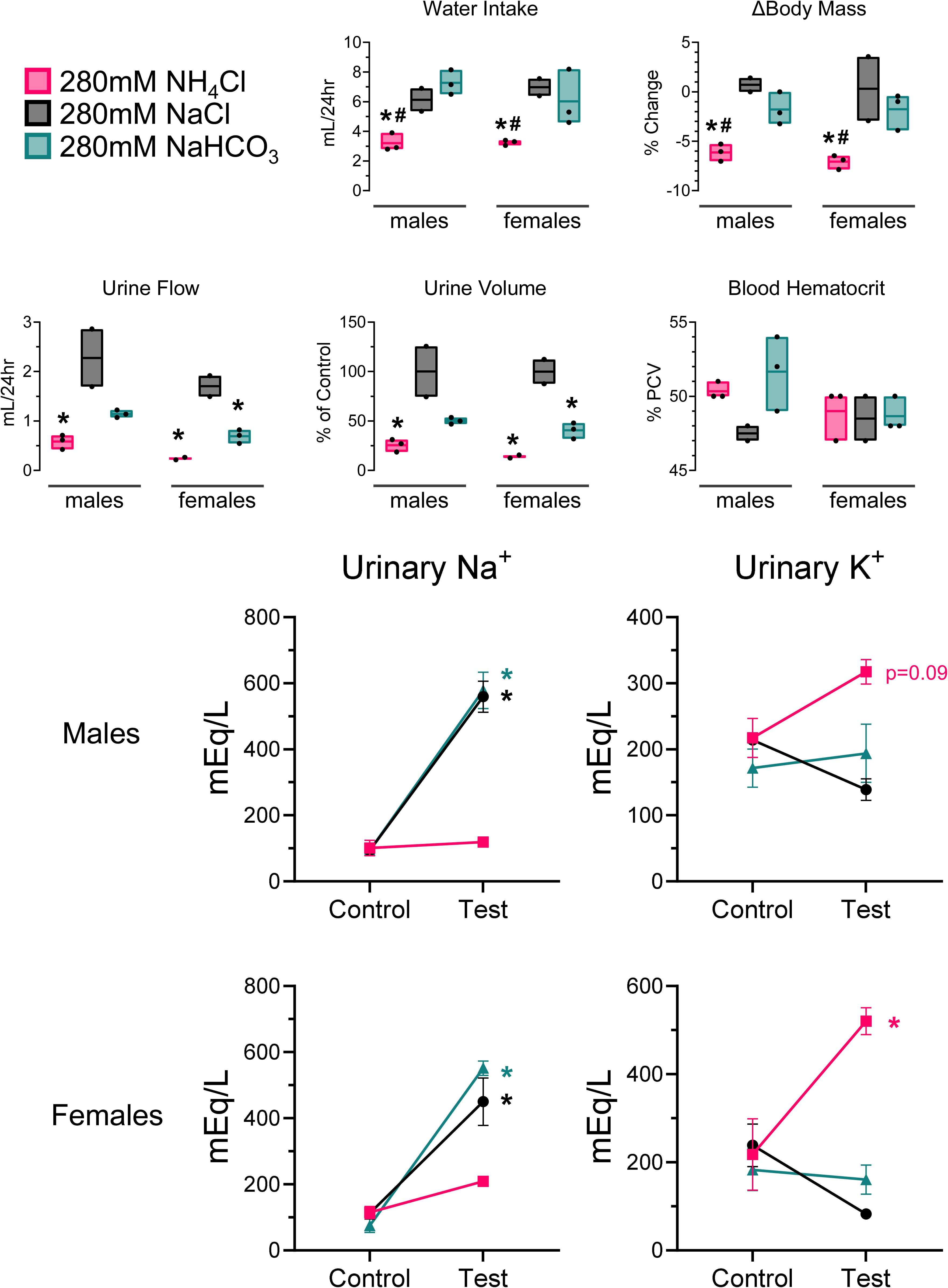
Physiological assessment of mice following acute pH challenge. % PCV = packed cell volume. * p_adj_<0.05 test vs control, # p_adj_<0.05 acidosis vs alkalosis.

**Supplemental Figure 2:**
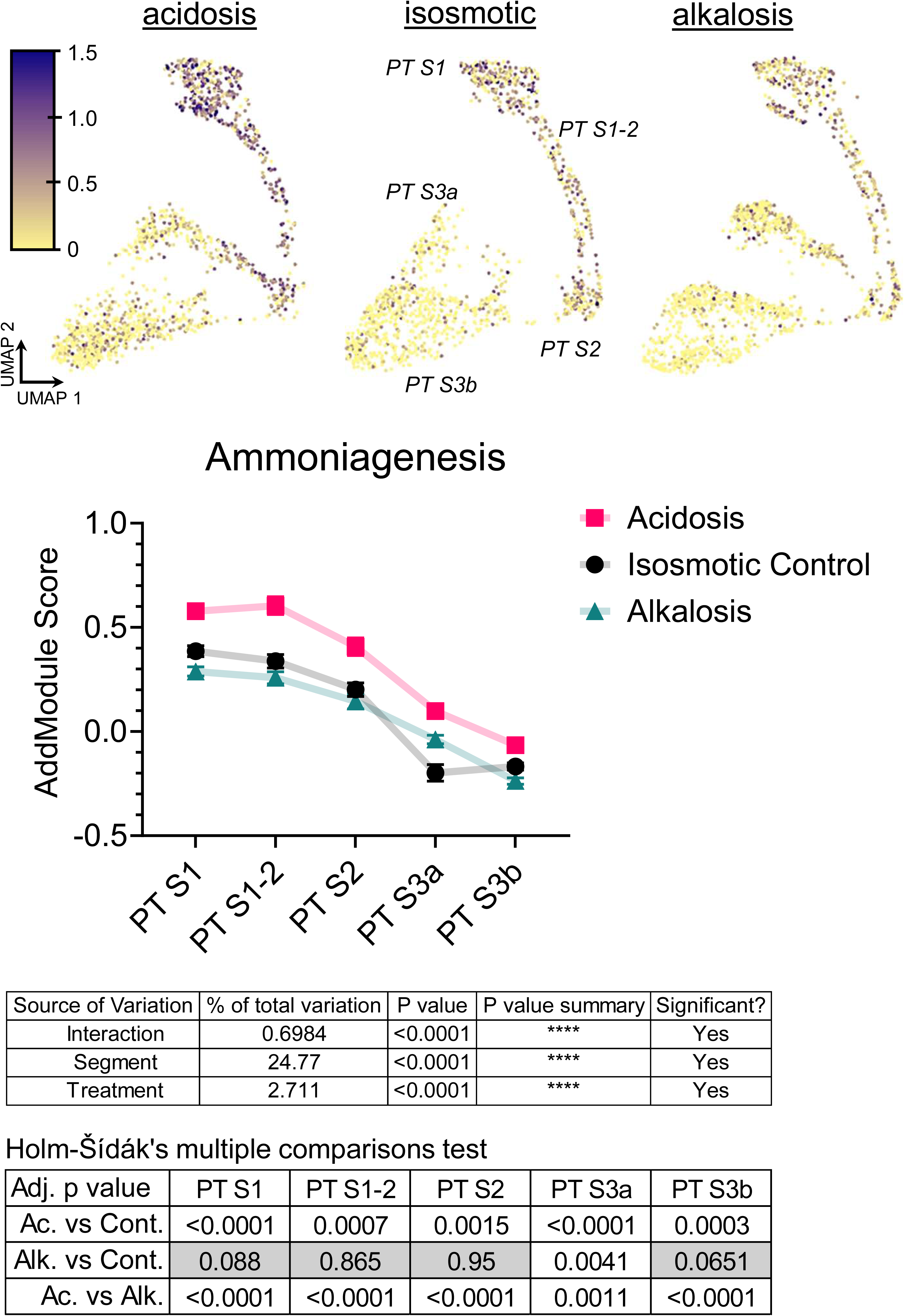
Quantification of male ammoniagenesis AddModule Score analysis. Data points are means ± SEM. Additional statistical analysis is available in the Supplemental Tables.

**Supplemental Figure 3:**
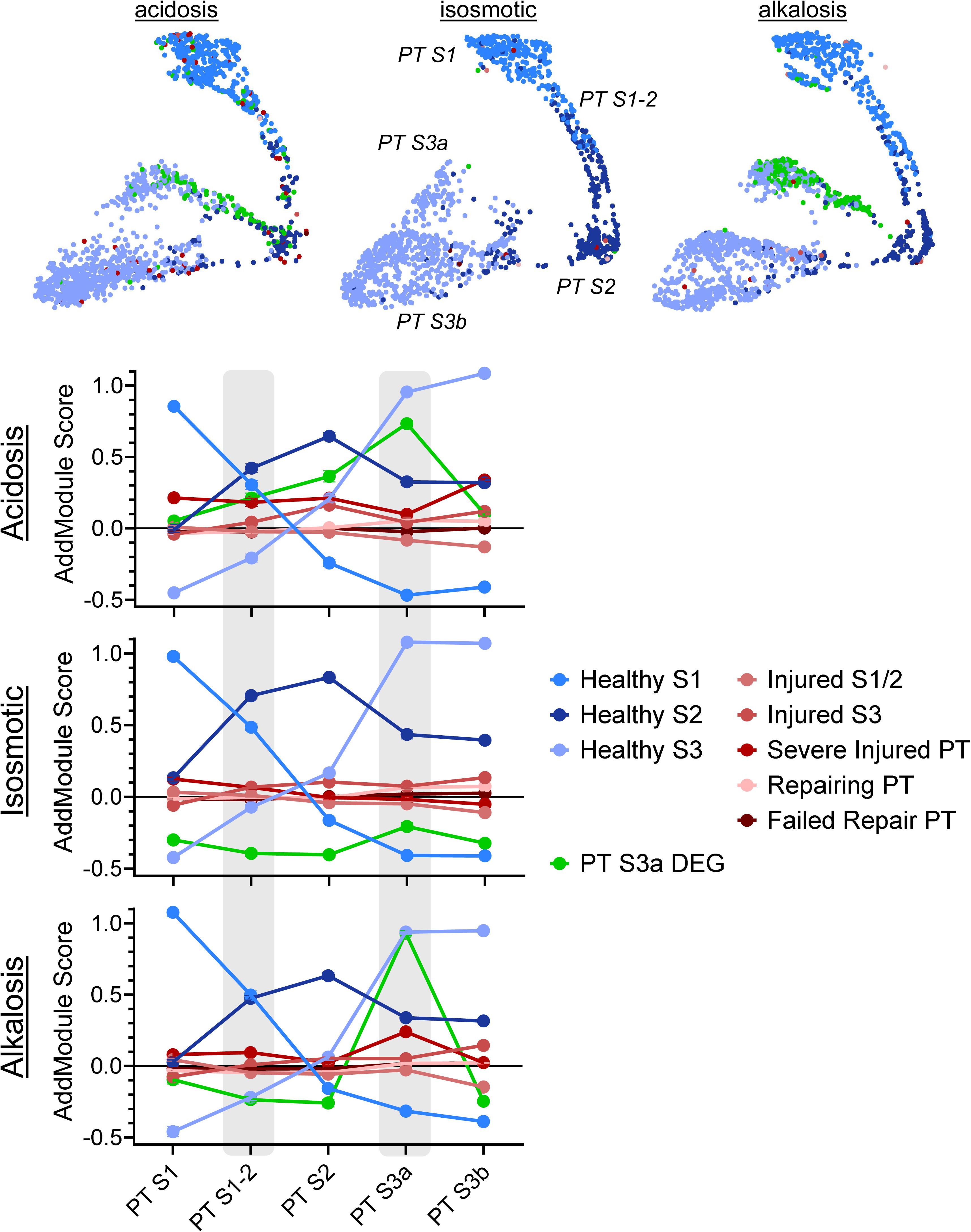
Quantification of male injury AddModule Score analysis. Data points are means ± SEM.

**Supplemental Figure 4:**
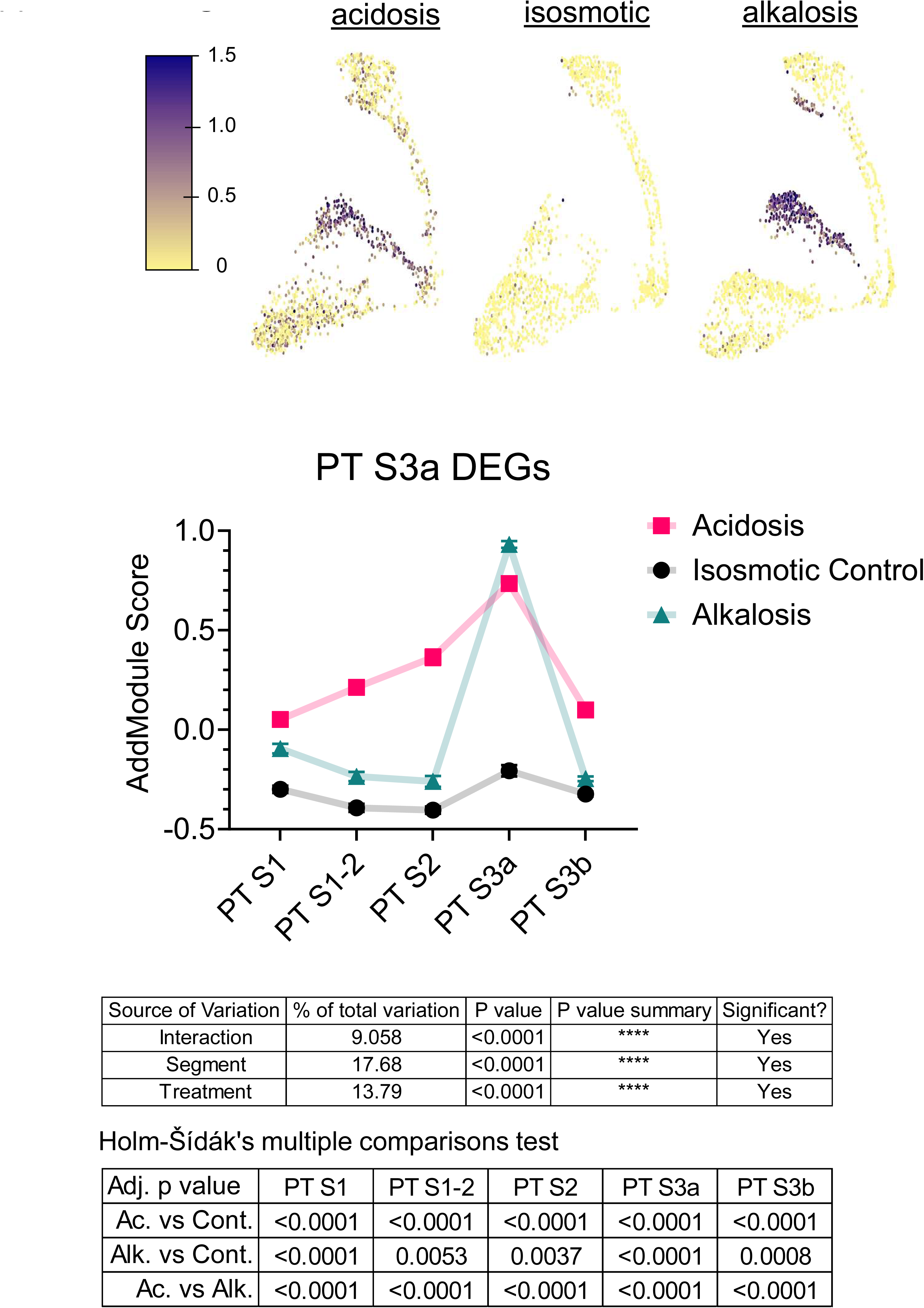
Quantification of male PT S3a DEGs AddModule Score analysis. Data points are means ± SEM. Additional statistical analysis is available in the Supplemental Tables.

**Supplemental Figure 5:**
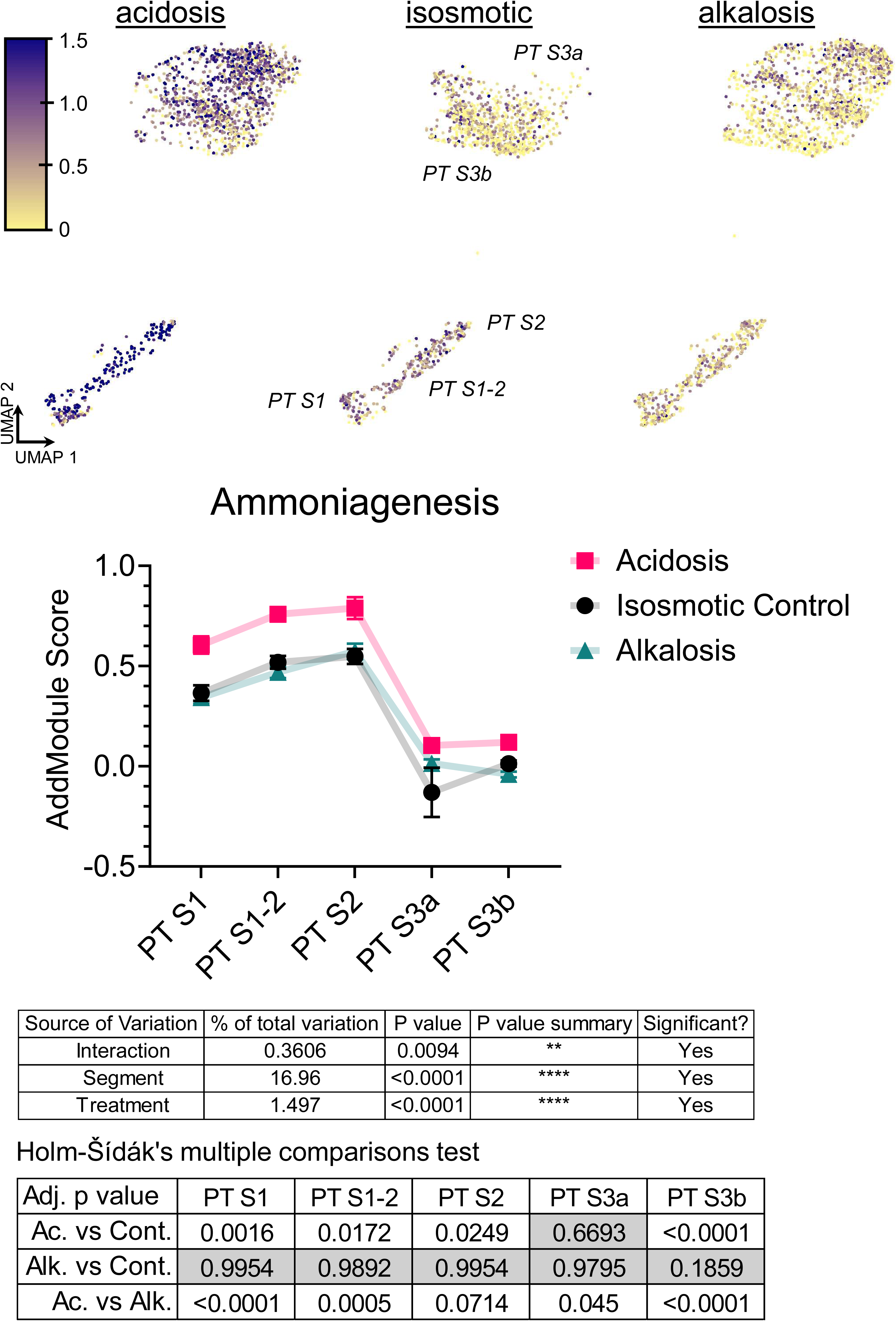
Quantification of female ammoniagenesis AddModule Score analysis. Data points are means ± SEM. Additional statistical analysis is available in the Supplemental Tables.

**Supplemental Figure 6:**
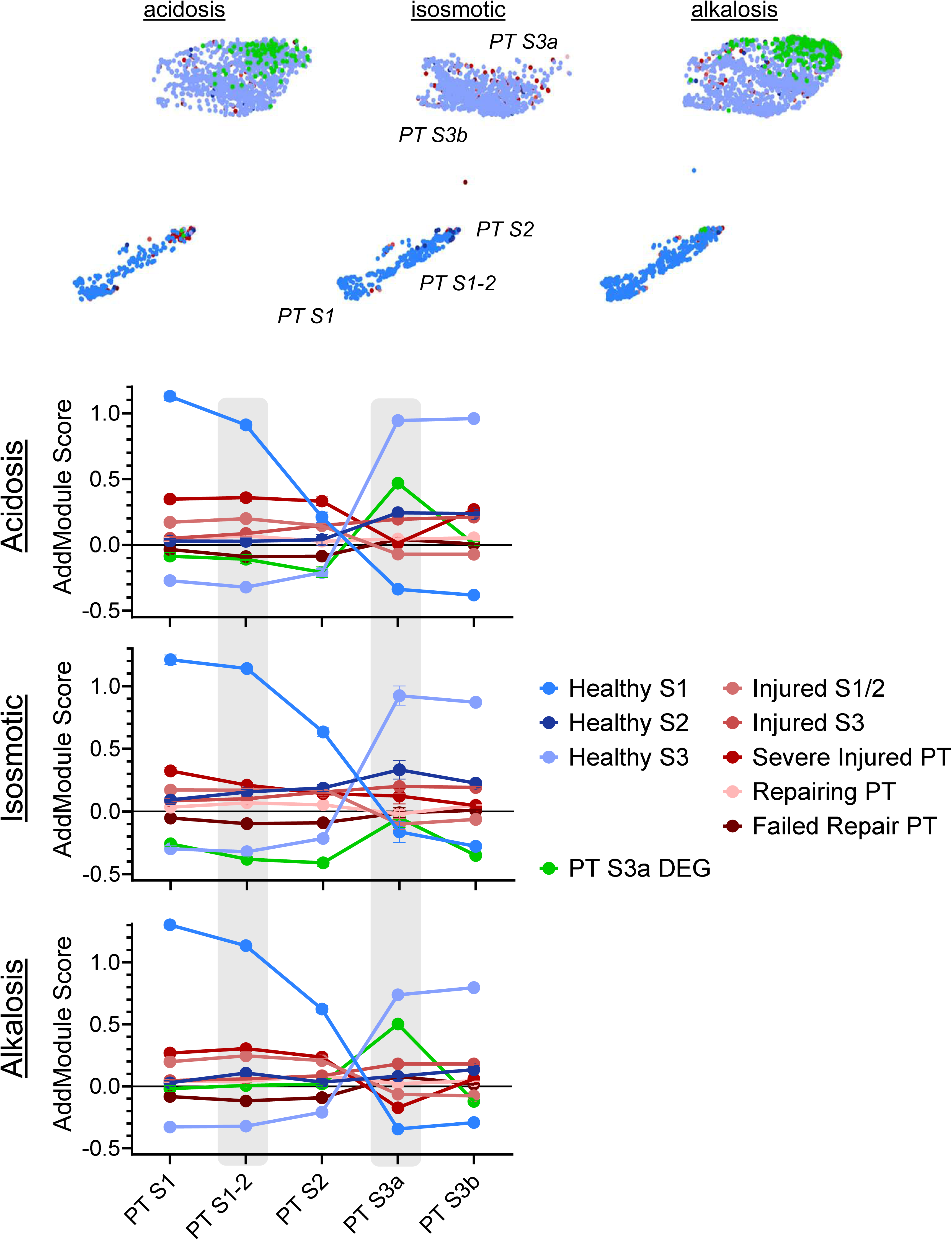
Quantification of female injury AddModule Score analysis. Data points are means ± SEM.

**Supplemental Figure 7:**
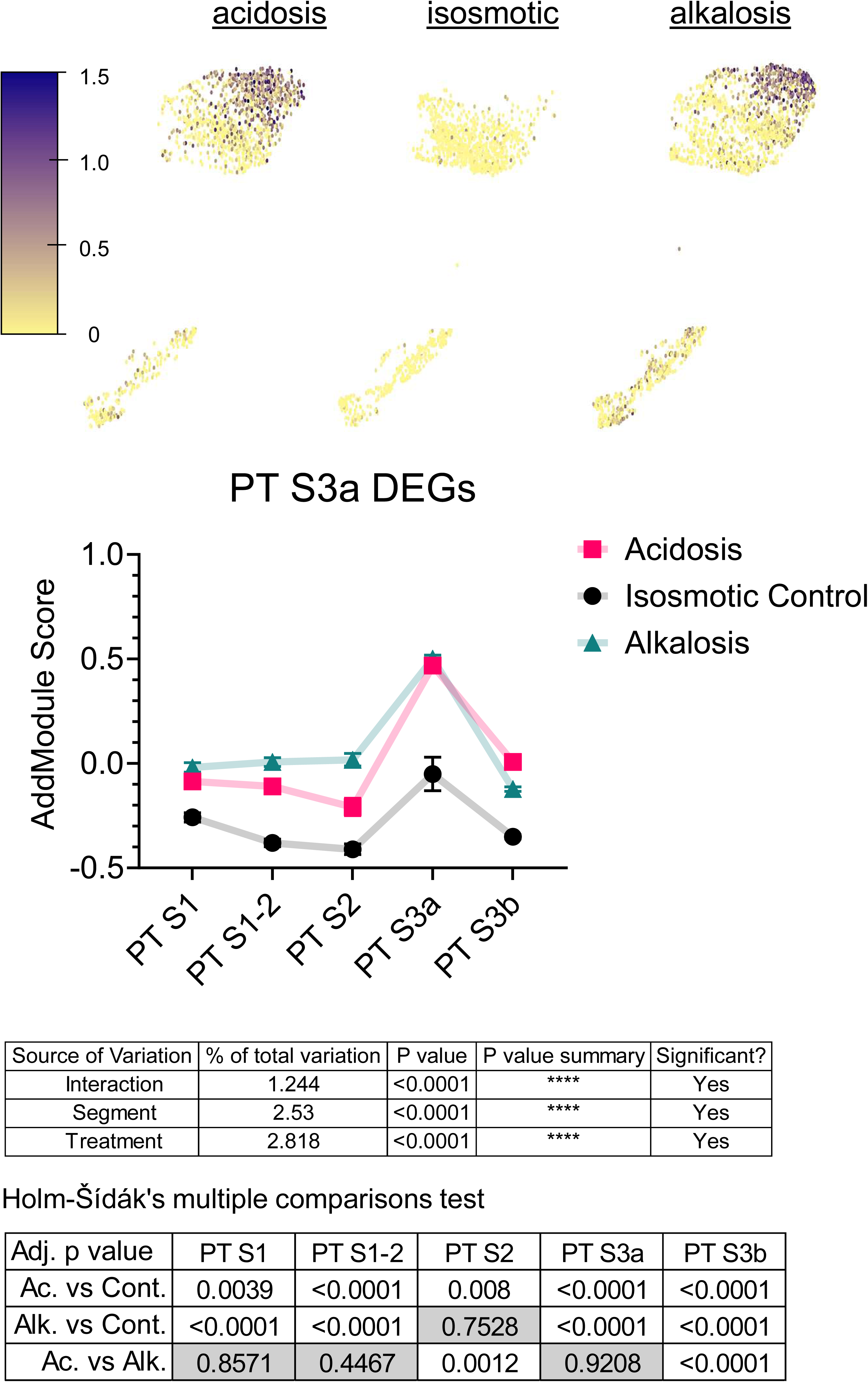
Quantification of female PT S3a DEGs AddModule Score analysis. Data points are means ± SEM. Additional statistical analysis is available in the Supplemental Tables.

**Supplemental Figure 8:**
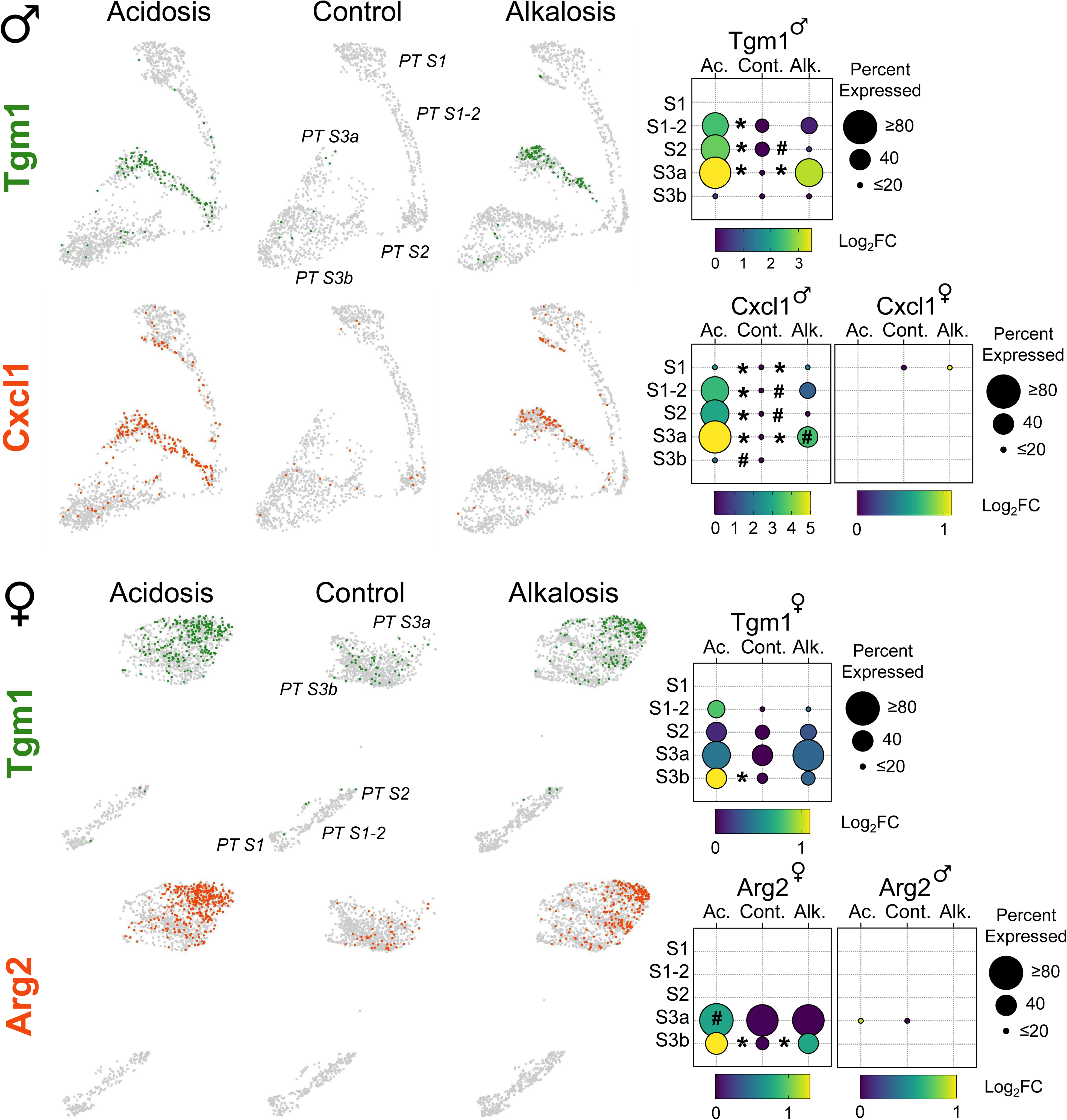
Sex-dependent expression of genes specific to PT S3a in male and female mice.

## References

1. Imenez Silva PH, Mohebbi N: Kidney metabolism and acid-base control: back to the basics. Pflugers Arch, 474: 919–934, 2022 10.1007/s00424-022-02696-6

2. Bushinsky DA, Krieger NS: Effects of acid on bone. Kidney Int, 101: 1160–1170, 2022 10.1016/j.kint.2022.02.032

3. Ho JQ, Abramowitz MK: Clinical Consequences of Metabolic Acidosis-Muscle. Adv Chronic Kidney Dis, 29: 395–405, 2022 10.1053/j.ackd.2022.04.010

4. Lee Hamm L, Hering-Smith KS, Nakhoul NL: Acid-base and potassium homeostasis. Semin Nephrol, 33: 257–264, 2013 10.1016/j.semnephrol.2013.04.006

5. Urso C, Brucculeri S, Caimi G: Acid-base and electrolyte abnormalities in heart failure: pathophysiology and implications. Heart Fail Rev, 20: 493–503, 2015 10.1007/s10741-015-9482-y

6. Urso C, Canino B, Brucculeri S, Firenze A, Caimi G: Analysis of electrolyte abnormalities and the mechanisms leading to arrhythmias in heart failure. A literature review. Clin Ter, 167: e85–91, 2016 10.7417/CT.2016.1944

7. Wagner CA, Unwin R, Lopez-Garcia SC, Kleta R, Bockenhauer D, Walsh S: The pathophysiology of distal renal tubular acidosis. Nat Rev Nephrol, 19: 384–400, 2023 10.1038/s41581-023-00699-9

8. Achanti A, Szerlip HM: Acid-Base Disorders in the Critically Ill Patient. Clin J Am Soc Nephrol, 18: 102–112, 2023 10.2215/CJN.04500422

9. Madias NE: Metabolic Acidosis and CKD Progression. Clin J Am Soc Nephrol, 16: 310-312, 2021 10.2215/CJN.07990520

10. Kim HJ: Metabolic Acidosis in Chronic Kidney Disease: Pathogenesis, Clinical Consequences, and Treatment. Electrolyte Blood Press, 19: 29-37, 2021 10.5049/EBP.2021.19.2.29

11. Melamed ML, Raphael KL: Metabolic Acidosis in CKD: A Review of Recent Findings. Kidney Med, 3: 267–277, 2021 10.1016/j.xkme.2020.12.006

12. Weiner ID, Verlander JW: Ammonia Transporters and Their Role in Acid-Base Balance. Physiol Rev, 97: 465–494, 2017 10.1152/physrev.00011.2016

13. Goraya N, Wesson DE: Pathophysiology of Diet-Induced Acid Stress. Int J Mol Sci, 25, 2024 10.3390/ijms25042336

14. Scialla JJ, Anderson CA: Dietary acid load: a novel nutritional target in chronic kidney disease? Adv Chronic Kidney Dis, 20: 141–149, 2013 10.1053/j.ackd.2012.11.001

15. Adam W, Simpson DP: Glutamine transport in rat kidney mitochondria in metabolic acidosis. J Clin Invest, 54: 165–174, 1974 10.1172/JCI107738

16. Alleyne GA, Scullard GH: Renal metabolic response to acid base changes. I. Enzymatic control of ammoniagenesis in the rat. J Clin Invest, 48: 364–370, 1969 10.1172/JCI105993

17. Deferrari G, Robaudo C, Garibotto G, Saffioti S, Sala MR, Tizianello A: Determinants of the partition of renal ammonia production between urine and venous blood in man with metabolic acid-base disturbances. Contrib Nephrol, 92: 109–113, 1991 10.1159/000420085

18. Nowik M, Lecca MR, Velic A, Rehrauer H, Brandli AW, Wagner CA: Genome-wide gene expression profiling reveals renal genes regulated during metabolic acidosis. Physiol Genomics, 32: 322–334, 2008 10.1152/physiolgenomics.00160.2007

19. Alleyne GA: Renal metabolic response to acid-base changes. II. The early effects of metabolic acidosis on renal metabolism in the rat. J Clin Invest, 49: 943-951, 1970 10.1172/JCI106314

20. Curthoys NP, Godfrey SS: Properties of rat kidney glutaminase enzymes and their role in renal ammoniagenesis. Curr Probl Clin Biochem, 6: 346–356, 1976

21. Amlal H, Paillard M, Bichara M: NH4+ transport pathways in cells of medullary thick ascending limb of rat kidney. NH4+ conductance and K+/NH4+(H+) antiport. J Biol Chem, 269: 21962-21971, 1994

22. Attmane-Elakeb A, Amlal H, Bichara M: Ammonium carriers in medullary thick ascending limb. Am J Physiol Renal Physiol, 280: F1–9, 2001 10.1152/ajprenal.2001.280.1.F1

23. Blanchard A, Eladari D, Leviel F, Tsimaratos M, Paillard M, Podevin RA: NH4+ as a substrate for apical and basolateral Na(+)-H+ exchangers of thick ascending limbs of rat kidney: evidence from isolated membranes. J Physiol, 506 ( Pt 3): 689–698, 1998 10.1111/j.1469-7793.1998.689bv.x

24. Flessner MF, Mejia R, Knepper MA: Ammonium and bicarbonate transport in isolated perfused rodent long-loop thin descending limbs. Am J Physiol, 264: F388–396, 1993 10.1152/ajprenal.1993.264.3.F388

25. Weiner ID, Verlander JW: Emerging Features of Ammonia Metabolism and Transport in Acid-Base Balance. Semin Nephrol, 39: 394–405, 2019 10.1016/j.semnephrol.2019.04.008

26. Caner T, Abdulnour-Nakhoul S, Brown K, Islam MT, Hamm LL, Nakhoul NL: Mechanisms of ammonia and ammonium transport by rhesus-associated glycoproteins. Am J Physiol Cell Physiol, 309: C747–758, 2015 10.1152/ajpcell.00085.2015

27. Bourgeois S, Bounoure L, Christensen EI, Ramakrishnan SK, Houillier P, Devuyst O, et al.: Haploinsufficiency of the ammonia transporter Rhcg predisposes to chronic acidosis: Rhcg is critical for apical and basolateral ammonia transport in the mouse collecting duct. J Biol Chem, 288: 5518–5529, 2013 10.1074/jbc.M112.441782

28. Bastani B, Gluck S: Adaptational changes in renal vacuolar H(+)-ATPase in the rat remnant kidney. J Am Soc Nephrol, 8: 868–879, 1997 10.1681/ASN.V86868

29. Frank AE, Wingo CS, Andrews PM, Ageloff S, Knepper MA, Weiner ID: Mechanisms through which ammonia regulates cortical collecting duct net proton secretion. Am J Physiol Renal Physiol, 282: F1120–1128, 2002 10.1152/ajprenal.00266.2001

30. Kirita Y, Wu H, Uchimura K, Wilson PC, Humphreys BD: Cell profiling of mouse acute kidney injury reveals conserved cellular responses to injury. Proc Natl Acad Sci U S A, 117: 15874–15883, 2020 10.1073/pnas.2005477117

31. Wilson PC, Wu H, Kirita Y, Uchimura K, Ledru N, Rennke HG, et al.: The single-cell transcriptomic landscape of early human diabetic nephropathy. Proc Natl Acad Sci U S A, 116: 19619–19625, 2019 10.1073/pnas.1908706116

32. Muto Y, Yoshimura Y, Wu H, Chang-Panesso M, Ledru N, Woodward OM, et al.: Multiomics profiling of mouse polycystic kidney disease progression at a single-cell resolution. Proc Natl Acad Sci U S A, 121: e2410830121, 2024 10.1073/pnas.2410830121

33. Abedini A, Levinsohn J, Klotzer KA, Dumoulin B, Ma Z, Frederick J, et al.: Single-cell multi-omic and spatial profiling of human kidneys implicates the fibrotic microenvironment in kidney disease progression. Nat Genet, 56: 1712–1724, 2024 10.1038/s41588-024-01802-x

34. Wu H, Gonzalez Villalobos R, Yao X, Reilly D, Chen T, Rankin M, et al.: Mapping the single-cell transcriptomic response of murine diabetic kidney disease to therapies. Cell Metab, 34: 1064–1078 e1066, 2022 10.1016/j.cmet.2022.05.010

35. Abedini A, Sanchez-Navaro A, Wu J, Klotzer KA, Ma Z, Poudel B, et al.: Single-cell transcriptomics and chromatin accessibility profiling elucidate the kidney-protective mechanism of mineralocorticoid receptor antagonists. J Clin Invest, 134, 2024 10.1172/JCI157165

36. Gerhardt LMS, Koppitch K, van Gestel J, Guo J, Cho S, Wu H, et al.: Lineage Tracing and Single-Nucleus Multiomics Reveal Novel Features of Adaptive and Maladaptive Repair after Acute Kidney Injury. J Am Soc Nephrol, 34: 554–571, 2023 10.1681/ASN.0000000000000057

37. Wu H, Kirita Y, Donnelly EL, Humphreys BD: Advantages of Single-Nucleus over Single-Cell RNA Sequencing of Adult Kidney: Rare Cell Types and Novel Cell States Revealed in Fibrosis. J Am Soc Nephrol, 30: 23–32, 2019 10.1681/ASN.2018090912

38. Zheng GX, Terry JM, Belgrader P, Ryvkin P, Bent ZW, Wilson R, et al.: Massively parallel digital transcriptional profiling of single cells. Nat Commun, 8: 14049, 2017 10.1038/ncomms14049

39. Stuart T, Butler A, Hoffman P, Hafemeister C, Papalexi E, Mauck WM, 3rd, et al.: Comprehensive Integration of Single-Cell Data. Cell, 177: 1888–1902 e1821, 2019 10.1016/j.cell.2019.05.031

40. Satija R, Farrell JA, Gennert D, Schier AF, Regev A: Spatial reconstruction of single-cell gene expression data. Nat Biotechnol, 33: 495–502, 2015 10.1038/nbt.3192

41. Hafemeister C, Satija R: Normalization and variance stabilization of single-cell RNA-seq data using regularized negative binomial regression. Genome Biol, 20: 296, 2019 10.1186/s13059-019-1874-1

42. Cao J, Spielmann M, Qiu X, Huang X, Ibrahim DM, Hill AJ, et al.: The single-cell transcriptional landscape of mammalian organogenesis. Nature, 566: 496–502, 2019 10.1038/s41586-019-0969-x

43. Yu G, Wang LG, Han Y, He QY: clusterProfiler: an R package for comparing biological themes among gene clusters. OMICS, 16: 284–287, 2012 10.1089/omi.2011.0118

44. Subramanian A, Tamayo P, Mootha VK, Mukherjee S, Ebert BL, Gillette MA, et al.: Gene set enrichment analysis: a knowledge-based approach for interpreting genome-wide expression profiles. Proc Natl Acad Sci U S A, 102: 15545–15550, 2005 10.1073/pnas.0506580102

45. Liberzon A, Subramanian A, Pinchback R, Thorvaldsdottir H, Tamayo P, Mesirov JP: Molecular signatures database (MSigDB) 3.0. Bioinformatics, 27: 1739–1740, 2011 10.1093/bioinformatics/btr260

46. Liberzon A, Birger C, Thorvaldsdottir H, Ghandi M, Mesirov JP, Tamayo P: The Molecular Signatures Database (MSigDB) hallmark gene set collection. Cell Syst, 1: 417–425, 2015 10.1016/j.cels.2015.12.004

47. Castanza AS, Recla JM, Eby D, Thorvaldsdottir H, Bult CJ, Mesirov JP: Extending support for mouse data in the Molecular Signatures Database (MSigDB). Nat Methods, 20: 1619–1620, 2023 10.1038/s41592-023-02014-7

48. Ponomarova O, Starbard AN, Belfi A, Anderson AV, Sundaram MV, Walhout AJ: idh-1 neomorphic mutation confers sensitivity to vitamin B12 in Caenorhabditis elegans. Life Sci Alliance, 7, 2024 10.26508/lsa.202402924

49. Gee MT, Kurtz I, Pannabecker TL: Expression of SLC4A11 protein in mouse and rat medulla: a candidate transporter involved in outer medullary ammonia recycling. Physiol Rep, 7: e14089, 2019 10.14814/phy2.14089

50. Su XT, Reyes JV, Lackey AE, Demirci H, Bachmann S, Maeoka Y, et al.: Enriched Single-Nucleus RNA-Sequencing Reveals Unique Attributes of Distal Convoluted Tubule Cells. J Am Soc Nephrol, 35: 426–440, 2024 10.1681/ASN.0000000000000297

51. Nath KA, Hostetter MK, Hostetter TH: Pathophysiology of chronic tubulo-interstitial disease in rats. Interactions of dietary acid load, ammonia, and complement component C3. J Clin Invest, 76: 667-675, 1985 10.1172/JCI112020

52. Tolins JP, Hostetter MK, Hostetter TH: Hypokalemic nephropathy in the rat. Role of ammonia in chronic tubular injury. J Clin Invest, 79: 1447–1458, 1987 10.1172/JCI112973

53. Wesson DE, Simoni J: Acid retention during kidney failure induces endothelin and aldosterone production which lead to progressive GFR decline, a situation ameliorated by alkali diet. Kidney Int, 78: 1128–1135, 2010 10.1038/ki.2010.348

54. Nagami GT: Role of angiotensin II in the enhancement of ammonia production and secretion by the proximal tubule in metabolic acidosis. Am J Physiol Renal Physiol, 294: F874–880, 2008 10.1152/ajprenal.00286.2007

55. Wesson DE, Buysse JM, Bushinsky DA: Mechanisms of Metabolic Acidosis-Induced Kidney Injury in Chronic Kidney Disease. J Am Soc Nephrol, 31: 469–482, 2020 10.1681/ASN.2019070677

56. Chen JW, Huang MJ, Chen XN, Wu LL, Li QG, Hong Q, et al.: Transient upregulation of EGR1 signaling enhances kidney repair by activating SOX9(+) renal tubular cells. Theranostics, 12: 5434–5450, 2022 10.7150/thno.73426

57. Livingston MJ, Zhang M, Kwon SH, Chen JK, Li H, Manicassamy S, et al.: Autophagy activates EGR1 via MAPK/ERK to induce FGF2 in renal tubular cells for fibroblast activation and fibrosis during maladaptive kidney repair. Autophagy, 20: 1032–1053, 2024 10.1080/15548627.2023.2281156

58. Shi L, Zha H, Pan Z, Wang J, Xia Y, Li H, et al.: DUSP1 protects against ischemic acute kidney injury through stabilizing mtDNA via interaction with JNK. Cell Death Dis, 14: 724, 2023 10.1038/s41419-023-06247-4

59. Jiao Y, Liu X, Shi J, An J, Yu T, Zou G, et al.: Unraveling the interplay of ferroptosis and immune dysregulation in diabetic kidney disease: a comprehensive molecular analysis. Diabetol Metab Syndr, 16: 86, 2024 10.1186/s13098-024-01316-w

60. Shah SN, Abramowitz M, Hostetter TH, Melamed ML: Serum bicarbonate levels and the progression of kidney disease: a cohort study. Am J Kidney Dis, 54: 270–277, 2009 10.1053/j.ajkd.2009.02.014

61. Raphael KL, Wei G, Baird BC, Greene T, Beddhu S: Higher serum bicarbonate levels within the normal range are associated with better survival and renal outcomes in African Americans. Kidney Int, 79: 356–362, 2011 10.1038/ki.2010.388

62. Raphael KL, Zhang Y, Wei G, Greene T, Cheung AK, Beddhu S: Serum bicarbonate and mortality in adults in NHANES III. Nephrol Dial Transplant, 28: 1207–1213, 2013 10.1093/ndt/gfs609

63. Navaneethan SD, Schold JD, Arrigain S, Jolly SE, Wehbe E, Raina R, et al.: Serum bicarbonate and mortality in stage 3 and stage 4 chronic kidney disease. Clin J Am Soc Nephrol, 6: 2395–2402, 2011 10.2215/CJN.03730411

64. Dobre M, Yang W, Chen J, Drawz P, Hamm LL, Horwitz E, et al.: Association of serum bicarbonate with risk of renal and cardiovascular outcomes in CKD: a report from the Chronic Renal Insufficiency Cohort (CRIC) study. Am J Kidney Dis, 62: 670–678, 2013 10.1053/j.ajkd.2013.01.017

65. Khatir DS, Carlsen RK, Ivarsen P, Jespersen B, Pedersen M, Christensen KL, et al.: Effects of enhanced versus reduced vasodilating treatment on brachial and central blood pressure in patients with chronic kidney disease: a randomized controlled trial. J Hypertens, 39: 2232–2240, 2021 10.1097/HJH.0000000000002942

66. Carlsen RK, Khatir DS, Jensen D, Birn H, Buus NH: Prediction of CKD Progression and Cardiovascular Events Using Albuminuria and Pulse Wave Velocity. Kidney Blood Press Res, 48: 468–475, 2023 10.1159/000530887

67. Misella Hansen N, Kamper AL, Rix M, Feldt-Rasmussen B, Leipziger J, Sorensen MV, et al.: Health effects of the New Nordic Renal Diet in patients with stage 3 and 4 chronic kidney disease, compared with habitual diet: a randomized trial. Am J Clin Nutr, 118: 1042–1054, 2023 10.1016/j.ajcnut.2023.08.008

68. Svendsen SL, Rousing AQ, Carlsen RK, Khatir D, Jensen D, Hansen NM, et al.: A Urine pH-Ammonium Acid/Base Score and CKD Progression. J Am Soc Nephrol, 35: 1533–1545, 2024 10.1681/ASN.0000000000000447

69. Rhee CM, Kovesdy CP: Epidemiology: Spotlight on CKD deaths-increasing mortality worldwide. Nat Rev Nephrol, 11: 199–200, 2015 10.1038/nrneph.2015.25

70. Jager KJ, Kovesdy C, Langham R, Rosenberg M, Jha V, Zoccali C: A single number for advocacy and communication-worldwide more than 850 million individuals have kidney diseases. Kidney Int, 96: 1048–1050, 2019 10.1016/j.kint.2019.07.012

